# Twin-arginine transport complex plays an essential role in *Caulobacter* cell shape and viability

**DOI:** 10.1101/2025.10.31.685914

**Authors:** Trisha N. Chong, Klara Christensen, Damion L. Whitfield, Mayura Panjalingam, Nima Pendar, Phway Phway Myat, Joseph C. Chen

## Abstract

Two main pathways are responsible for protein secretion across the cytoplasmic membrane in prokaryotes. While the general secretory (Sec) pathway transports proteins across the membrane in an unfolded state, the twin-arginine translocation (Tat) pathway transports proteins primarily in their folded conformation. Although the Tat system appears dispensable in multiple model bacteria, some species require it for viability, and the reason for the distinction is nebulous. Here we show that all three subunits of the Tat complex -- TatA, TatB, and TatC -- are essential in the alpha-proteobacterium *Caulobacter crescentus*. Additionally, depletion of the Tat complex results in abnormal cell morphology. We found that localization to the cell periphery, as well as midcell localization upon osmotic upshift, of the essential peptidoglycan transpeptidase PBP2 is dependent on the Tat apparatus. In contrast, subcellular localization of the actin homolog MreB and the penicillin-binding protein PBP1a is not perturbed upon depletion of the Tat complex. PBP2 transpeptidase activity links glycan chains at sites of cell wall remodeling and is essential for cell elongation. Together these results suggest that PBP2 localization is a key responsibility of the Tat system in *Caulobacter* and possibly other alpha-proteobacteria.

**Significance Statement:**

- The twin-arginine translocation (Tat) system is essential for viability in some bacteria but not others. The essential role that the Tat pathway plays in these bacteria is not well understood.
- The Tat complex is essential in *Caulobacter crescentus* and required for the cell wall synthesis protein PBP2 to localize to the cell envelope.
- PBP2 is critical for viability and maintenance of cell shape in *Caulobacter*, and essentiality of the Tat complex may be partly attributed to its role in localizing PBP2.

## Introduction

Gram-negative bacteria have two main secretory pathways, Sec and Tat, that are responsible for exporting proteins across the cytoplasmic membrane and integrating the hydrophobic helices of membrane proteins into the lipid bilayer (Gallego-Parrilla *et al*., 2024; Hatzixanthis *et al*., 2003; Palmer & Stansfeld, 2020; Tooke *et al*., 2017). The vast majority of exported proteins are transported via the general secretory (Sec) pathway: they are threaded through the SecYEG translocon in an unfolded state and then folded in the extracellular compartment (Crane & Randall, 2017; Kuhn *et al*., 2017). In contrast, the twin-arginine translocation (Tat) system ferries proteins in a folded state, often those bound noncovalently to prosthetic groups, across the inner membrane (Natale *et al*., 2008; Palmer & Berks, 2024). Tat-secreted proteins have an N-terminal signal peptide distinct from Sec-secreted proteins in that they generally contain a twin-arginine motif (RR) in addition to a weakly hydrophobic region and a positively charged Sec-avoidance region (Frain *et al*., 2019; Palmer & Stansfeld, 2020; Pradel *et al*., 2009). Substrates of the Tat pathway contribute to diverse processes, including energy metabolism, cell wall biosynthesis, and virulence, and the number of substrates can vary greatly among species (Berks *et al*., 2005; De Buck *et al*., 2008; Dilks *et al*., 2003). While the Tat machinery is dispensable for viability in multiple model bacteria, including *Escherichia coli*, *Bacillus subtilis*, *Pseudomonas aeruginosa*, *Streptomyces coelicolor*, and *Yersinia pestis* and *Y. pseudotuberculosis* (Avican *et al*., 2016; Bozue *et al*., 2014; Jongbloed *et al*., 2002; Ochsner *et al*., 2002; Sargent *et al*., 1999; Voulhoux *et al*., 2001; Widdick *et al*., 2006), it appears to be essential in a few species, such as *Mycobacterium tuberculosis* and *Bdellavibrio bacteriovorous* (Chang *et al*., 2011; Palmer & Berks, 2012; Saint-Joanis *et al*., 2006). Why the Tat system is essential in some bacteria but not others has not been explored in depth: for example, for two alpha-proteobacteria in the *Rhizobiaceae* family, the Tat complex is essential in *Sinorhizobium meliloti* but dispensable in *Agrobacterium tumefaciens* (Ding & Christie, 2003; Pickering & Oresnik, 2010). In *Caulobacter crescentus*, another alpha-proteobacterium, previous transposon insertion screens indicated that all three subunits that comprise the Tat complex (TatA, TatB, and TatC) are essential (Christen *et al*., 2011; Price *et al*., 2018), suggesting that one or more Tat-dependent substrates is critical for *Caulobacter* survival and proliferation.

Various substrates of the Tat pathway in *E. coli*, such as the AmiA and AmiC amidases, and in other bacteria participate in cell wall maintenance and metabolism (Bernhardt & de Boer, 2003; Brauer *et al*., 2021; Gallego-Parrilla *et al*., 2024; Ize *et al*., 2003; Saint-Joanis *et al*., 2006). The bacterial cell wall is essential for growth and viability, as it gives cells their defined shape and prevents osmotic lysis (Höltje, 1998). In most bacteria, this cell wall or peptidoglycan (PG) layer is composed of glycan chains cross-linked by short peptides. Throughout growth and division, the PG layer must be remodeled to permit changes in cell shape (Sauvage *et al*., 2008; Spratt, 1975). Studies in *E. coli* and other species indicated that PG remodeling during cell elongation is carried out by the Rod complex (Garde *et al*., 2021). Main components of the Rod complex include the PG polymerase RodA and the transpeptidase PBP2, which associate with the actin-like cytoskeletal protein MreB and multiple other proteins, including MreC, MreD, RodZ, and PBP1a; the cytoskeletal elements MreBCD form a dynamic scaffold that guides peptidoglycan synthesis by RodA and PBP2 (Banzhaf *et al*., 2012; Bratton *et al*., 2018; Liu *et al*., 2020; Morgenstein *et al*., 2015; Özbaykal *et al*., 2020; Rohs *et al*., 2018). For transpeptidase activity, PBP2 must be inserted into the inner membrane with its C-terminal enzymatic domain localized to the periplasm (Nygaard *et al*., 2023; Shlosman *et al*., 2023). Some evidence suggests that PBP2 translocation is carried out by the Sec translocase and/or YidC insertase in *E. coli* (de Sousa Borges *et al*., 2015). In contrast, *Caulobacter* PBP2 contains an N-terminal signal peptide that is consistent with Tat-mediated translocation (Dilks *et al*., 2003; Teufel *et al*., 2022).

Here, we present evidence that the Tat system and PBP2 are both essential for viability and maintenance of cell shape in *Caulobacter*. Moreover, as the subcellular localization of PBP2 depends on the Tat complex, we propose that PBP2 translocation is a vital function of the Tat complex in *Caulobacter*.

## Results and Discussion

### Tat complex and PBP2 are essential for *Caulobacter* proliferation

While the Tat complex is dispensable in *E. coli,* all three components of the Tat complex were identified as essential genes by Tn-seq analyses of the *Caulobacter* genome (Christen *et al*., 2011; Price *et al*., 2018). To confirm the essentiality of the Tat complex proteins, we used a plasmid loss assay (Arellano *et al*., 2010) to determine whether a plasmid harboring a *tat* gene could be lost by the same *tat* mutant: such plasmids carry *tatA, tatB,* or *tatC* along with an *E. coli lacZ* gene and a chloramphenicol resistance cassette. Cells that retained the plasmid formed blue colonies when grown on solid media containing 5-bromo-4-chloro-3-indolyl-beta-D-galactopyranoside (X-Gal) due to β-galactosidase activity conferred by *lacZ*. We observed that wild-type (WT) *Caulobacter* cells readily lost the *tatC* plasmid when grown on plates without chloramphenicol, giving rise to sectored and white colonies (Figure 1A, panel i). In contrast, all colonies formed by a mutant with in-frame deletion of the native *tatC* locus (Δ*tatC*) retained the blue color, indicating that the complementing plasmid was not lost in these colonies and that *tatC* is necessary for cell propagation (Figure 1A, panel ii). Addition of chloramphenicol to the plates resulted in plasmid retention by both WT and Δ*tatC* strains (Figure 1A, panels iii and iv). These results were recapitulated in Δ*tatA* and Δ*tatB* mutants complemented with plasmids harboring *tatA* and *tatB* genes, respectively (Figure S1A). Additionally, a mutant in which all three *tat* genes were deleted (Δ*tatABC*) was not able to lose a plasmid carrying the three genes (Figure S1A). We obtained similar results with both NA1000- and CB15-derived strains (Figure S1A).

**Figure 1.**
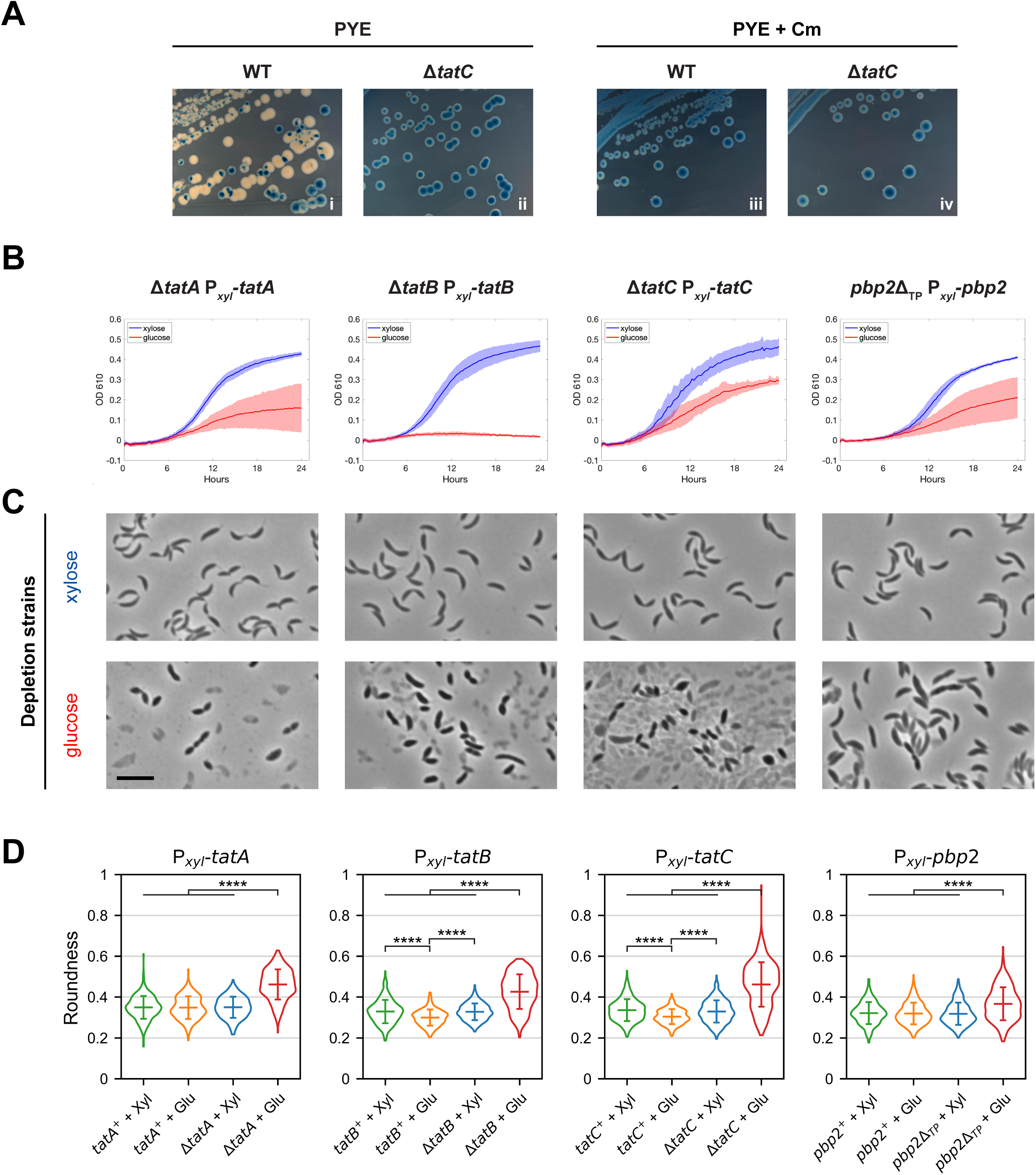
The Tat protein complex and PBP2 are critical for *Caulobacter* viability and cell shape. (**A**) Assessment of essentiality by plasmid loss. Wild-type (WT) and Δ*tatC* cells transformed with a plasmid harboring *tatC,* the *E. coli lacZ* gene, and a chloramphenicol resistance cassette were grown on PYE plates containing X-Gal, either with or without chloramphenicol (Cm). WT colonies grown without Cm frequently appeared white, indicating loss of the plasmid and *lacZ*. In contrast, all *ΔtatC* colonies turned blue, with or without Cm. (**B**) Growth of *tat* and *pbp2* strains. Depletion strains contain in-frame deletions at the native loci of individual *tat* genes or the region encoding the transpeptidase domain of PBP2, while complete sequences of corresponding genes were inserted downstream of the inducible *xylX* promoter (P*_xyl_*). Strains were grown in the presence of either 0.1% xylose (blue) or 0.1% glucose (red) for 24 hours, with OD610 measurements taken at 15-minute intervals. Lines indicate average readings, while shaded areas indicate standard deviations, derived from two independent trials, each with four technical replicates. (**C**) Cell morphology of Tat and PBP2 depletion strains. Phase contrast images show cells grown under inducing (xylose) or non-inducing (glucose) conditions for 22-24 hours, corresponding to those of the growth curves displayed above in (B). Scale bar, 5 µm. (**D**) Changes in cell morphology, as indicated by roundness. Expression strains (WT cells carrying P*_xyl_*transcriptional fusions) and depletion strains were grown with xylose (Xyl) or glucose (Glu) as above, and the roundness of cells in each population was measured. Violin plots depict distributions of the measurements, with horizontal bars representing means and standard deviations. Number of cells measured ranged from 246 to 400 per population. ****, *p* < 0.0001, based on two-tailed *t*-test. Images are representatives from at least two biological replicates.

To further verify the essentiality of the Tat complex, we generated depletion strains with deletions of each *tat* gene, complemented by a copy of the corresponding *tat* gene under the control of the endogenous, inducible *xylX* promoter (P*_xyl_*) on the chromosome (Stephens *et al*., 2007). Growth assays in PYE rich medium, with 0.1% xylose to induce expression of the complementing *tat* genes, showed that Tat depletion strains diluted to an initial optical density at 610 nm (OD610) of 0.01 reached final average OD610 of 0.43 - 0.46 after 24 hours of continuous shaking in 96-well plates (Figure 1B, blue curves). In contrast, cultures cultivated with 0.1% glucose to repress expression of *tatA, tatB,* or *tatC* exhibited reduction in growth, reaching average final OD610 of 0.16±0.12, 0.02±0.01, or 0.29±0.02, respectively (Figure 1B, red curves), suggesting that all *tat* genes are necessary for normal *Caulobacter* proliferation under these conditions. Variations in the timing and extent of growth arrests among the distinct Tat depletion strains may be due to differences in expression and cellular requirements of individual proteins. For instance, the TatB depletion strain appeared to stop doubling more quickly than the other strains after inoculation into PYE with glucose, possibly because TatB expression from P*_xyl_* is barely sufficient for normal operations, whereas TatA and TatC need more cell doublings for their protein levels to dip below the viable concentration requirements.

Next, we used strains carrying the P*_xyl_*-*tat* expression constructs to assess the influence of the Tat complex on cell morphology, by capturing phase contrast images of cells grown as above, after each *tat* gene had been induced or repressed for 22 - 24 hours. WT strains with the P*_xyl_*-*tat* constructs (expression strains) grown with xylose or glucose yielded typical crescent-shaped cells (Figure S1B), as did Tat depletion strains grown with xylose (Figure 1C). As for depletion strains grown with glucose, most cells instead exhibited a bloated, lemon-shaped morphology; moreover, significant fractions of cells appeared ruptured, ranging from approximately 30% in Δ*tatB* to 90% in Δ*tatA* and Δ*tatC* strains (Figure 1C). We therefore considered whether a protein that contributes to maintenance of cell shape may be impacted by the Tat complex. Because PBP2 transpeptidase activity is critical to PG remodeling, and it has a Tat recognition sequence in its N-terminus (Dilks *et al*., 2003; Teufel *et al*., 2022), it emerged as a candidate Tat substrate for further investigation.

To assess whether loss of PBP2 activity mimics loss of Tat activity, we first constructed a PBP2 depletion strain, with a mutant *pbp2* allele (*pbp2*Δ_TP_) that fused its R267 codon to L621, thus deleting the endogenous region encoding the transpeptidase domain, and intact *pbp2* under the control of the *xylX* promoter (P*_xyl_*-*pbp2*). As in another study (Wagner *et al*., 2005), we were unable to generate a deletion of the complete *pbp2* coding sequence, even while expressing it from the P*_xyl_*promoter, possibly due to polar effects on the essential *rodA* gene downstream (Christen *et al*., 2011). Growth assays with this depletion strain showed that cultures grown in the presence of 0.1% xylose or glucose reached an average OD610 of 0.41±0.01 or 0.21±0.10, respectively, after 24 hours, suggesting that *pbp2* activity is vital to *Caulobacter*’s proliferation under these conditions (Figure 1B), consistent with previously published reports (Christen *et al*., 2011; Price *et al*., 2018). We confirmed that PBP2 depletion strains had reduced growth on PYE plates supplemented with 0.1% glucose compared to both (1) the P*_xyl_*-*pbp2* expression strain and (2) the expression strain with the allelic replacement plasmid integrated into the genome, prior to deletion of *pbp2*’s transpeptidase region by homologous recombination (Figure S2A). This growth defect was rescued on PYE plates supplemented with 0.1% xylose (Figure S2A).

In addition, PBP2-depleted cells lost their crescent shape and became bloated (Figure 1C), akin to that seen previously with RodA-depleted *Caulobacter* cells (Wagner *et al*., 2005) and consistent with PBP2 and RodA forming the elongasome complex and participating in the same PG synthesis pathway, as shown in *E. coli* and *B. subtilis* (Cho *et al*., 2016; Meeske *et al*., 2016; Nygaard *et al*., 2023; Rohs *et al*., 2018; Shlosman *et al*., 2023). This change in morphology was similar, though not as severe, as that observed for Tat-depleted cells, with fewer lysed cells (<5%) in the PBP2-depleted population. MicrobeJ analysis (Ducret *et al*., 2016) indicated that depletion of individual Tat subunits or PBP2 led to significant increases in the average “roundness” of cells, as measured by 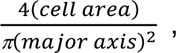 and the increase was smaller in PBP2-depleted cells compared to Tat-depleted cells (Figure 1D). Less extreme, albeit significant, differences in roundness were also observed when comparing TatB or TatC expression strains grown with glucose against their corresponding expression or depletion strains grown with xylose (Figure 1D). This small increase in roundness when TatB or TatC was ectopically expressed in both WT and deletion backgrounds seemed inconspicuous during visual inspection of the cell images; nevertheless, it may be that aberrant Tat complex stoichiometry perturbs normal protein translocation and morphology maintenance.

As for the difference in phenotype severity between Tat and PBP2 depletion strains, one possible explanation is that the PBP2 depletion strain may have more leaky expression of its complementing allele compared to the Tat depletion strains and thus less severe defects. Another possibility is that Tat substrates other than PBP2 also participate in the maintenance of cell shape, and thus disruption of the Tat pathway would lead to a more severe change in morphology. Finally, we cannot rule out that the Tat complex itself contributes to structural integrity of the cell envelope. Regardless of variations in growth and morphological defects among depletion strains, these results together indicate that all three components of the Tat complex and the transpeptidase function of PBP2 are essential for normal *Caulobacter* growth and cell shape, in agreement with previous studies that identified essential *Caulobacter* genes through Tn-seq analysis (Christen *et al*., 2011; Price *et al*., 2018).

### Tat complex is required for PBP2 midcell localization upon osmotic upshift

As PBP2-depleted cells had a similar phenotype to Tat-depleted cells, we asked if the Tat complex contributes to subcellular localization of PBP2. Hocking *et al*. showed that some membrane-bound proteins, including PBP2, accumulate quickly at the midcell during osmotic upshifts (Hocking *et al*., 2012). This shuttling to the midcell likely depends on proper insertion into the membrane. Thus, we utilized this observation to assess the effects of Tat depletion on PBP2, by imaging cells expressing *mCherry-pbp2* under the control of the chromosomal, vanillate-inducible *vanAB* promoter (P*_van_*) (Thanbichler *et al*., 2007) that were grown in PYE liquid media and then transferred to agarose pads made with M2, a defined medium with higher osmolality than PYE. This P*_van_*-*mCherry-pbp2* allele was able to complement the *pbp2*Δ_TP_ mutation, even in the absence of induction (Figure S2B). As anticipated, in WT cells grown with vanillate, we observed that mCherry-PBP2 localized to the midcell region (Figure 2A). In the Δ*tatA* background, cells grown with xylose to induce *tatA* expression from the P*_xyl_*promoter also exhibited midcell localization of mCherry-PBP2, whereas cells grown with glucose for four hours to deplete TatA showed diffuse or patchy fluorescence throughout the cell body (Figure 2A). Demographs of normalized fluorescence intensities along the medial cell axis (the length of the cell) confirmed that most of the mCherry-PBP2 signal was confined to the midcell region in WT and TatA-replete cells, while the signal in TatA*-*depleted cells was diffuse (Figure 2A). We also measured the integrated fluorescence along cell length, by calculating the area under the curve (AUC) of the normalized medial fluorescence profile, and found that the average AUC of TatA-depleted cells was significantly greater than those of WT and TatA-replete cells, indicating distinct localization patterns of mCherry-PBP2 (Figure 2B). These results suggested that the Tat complex is necessary for PBP2 localization in *Caulobacter*.

**Figure 2.**
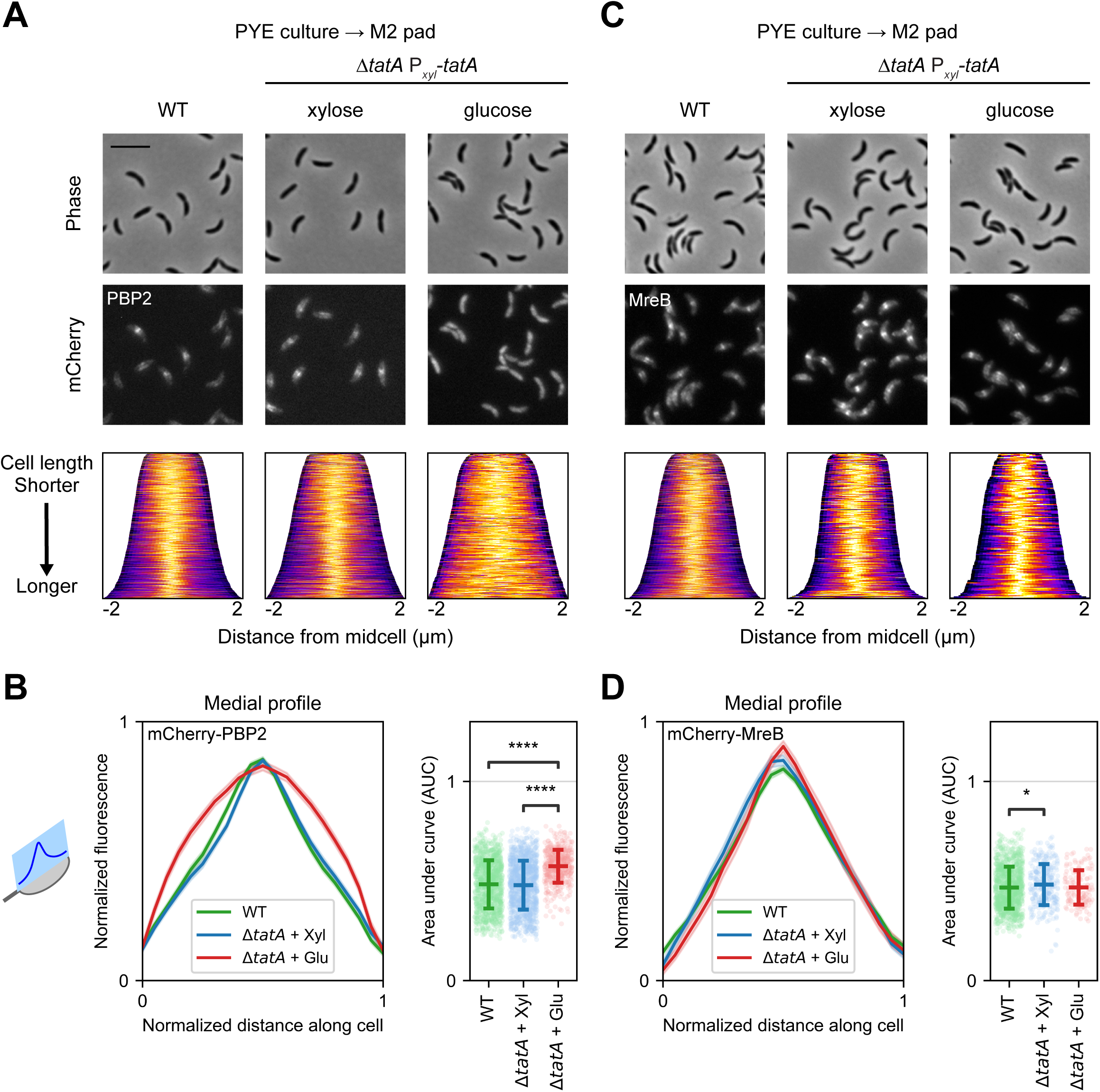
TatA is required for PBP2 midcell localization under osmotic shock. Cells were induced with vanillate to express (**A**, **B**) mCherry-PBP2 or (**C**, **D**) mCherry-MreB. TatA depletion strains (Δ*tatA* P*_xyl_*-*tatA*) were grown in the presence of xylose or glucose for 4 hours to express or repress *tatA*, respectively, and subjected to osmotic upshift from being transferred from PYE medium to an M2 agarose pad for microscopy. WT cells were treated similarly, except without glucose or xylose in the culture medium. (**A**, **C**) Representative phase contrast (top) and fluorescence (middle) images are shown with corresponding population-level demographs (bottom). Demographs depict localization of normalized fluorescence along the medial axis (the cell length), with cells arranged by length and lighter colors indicating brighter fluorescence. (**B**, **D**) Medial profiles (left panels) represent normalized fluorescence intensities along normalized cell length, as illustrated by the schematic of a model cell on the left. Colored lines indicate averages, while shaded areas indicate 95% confidence intervals. Area under curve was calculated for each medial profile and shown at the population level as scatter plots (right panels), with horizontal bars indicating means and standard deviations. (**A**, **B**) Midcell localization of mCherry-PBP2 depends on TatA; n = 1045 (WT), 1119 (Δ*tatA* + Xyl), 425 (Δ*tatA* + Glu). (**C**, **D**) Midcell localization of mCherry-MreB is not affected by TatA depletion; n = 1034 (WT), 134 (Δ*tatA* + Xyl), 291 (Δ*tatA* + Glu). Scale bar, 5 µm. *, *p* < 0.05; ****, *p* < 0.0001; based on two-tailed *t*-test.

We repeated osmotic upshift experiments with TatB and TatC depletion strains to evaluate the impacts of other Tat components on PBP2 localization. In the TatB depletion strains, we observed very limited midcell localization of mCherry-PBP2, whether in cells grown with xylose or glucose (Figure S3A). Nevertheless, calculations of AUC from normalized medial fluorescence profiles did indicate a small but significant difference between TatB-replete and TatB-depleted cells (Figure S3B). We posit three non-mutually exclusive explanations for these findings. First, expression of TatB from the P*_xyl_* promoter may be insufficient to allow optimal operation of the Tat system and efficient localization of mCherry-PBP2, particularly when mCherry-PBP2 is ectopically expressed. As the midcell localization signal is already low in TatB-replete cells, depletion of TatB leads to only a minute reduction in that signal. Another explanation is that the Δ*tatB* mutation affects expression of *tatC* downstream and could not be complemented fully by P*_xyl_*-*tatB* alone. Finally, suppressors may have arisen in the TatB depletion strains to allow survival despite poor Tat function and localization of PBP2.

As for the TatC depletion strain, cultivation with xylose to induce P*_xyl_*-*tatC* expression led to the same midcell localization of mCherry-PBP2 as WT and TatA-replete cells, but it required more time in PYE with glucose than the TatA depletion strain to yield mostly diffuse fluorescence: we observed clear loss of mCherry-PBP2 localization when *tatC* expression had been repressed for 20 hours (Figure S3, C and D), rather than four hours. These variations in the duration of depletion needed to alter the osmolarity-conditioned localization of mCherry-PBP2 may be attributed, again, to differences in the expression levels and cellular requirements of distinct Tat components, as mentioned above regarding growth arrests of depletion strains. Because both TatA and TatC depletion strains (as well as the TatB depletion strain, albeit less intuitively) indicated that PBP2 localization depends on the Tat complex in *Caulobacter*, we used primarily the TatA depletion strain for further analysis, as it allowed faster disruption of Tat functionality.

For comparison, we examined mCherry-MreB and mCherry-PBP1a localization in WT and TatA depletion strains. MreB localization to the midcell does not depend on osmotic upshift, while PBP1a does (Hocking *et al*., 2012). As expected, mCherry-MreB localized to the division site in WT and TatA-replete cells, and this pattern of localization was not perturbed in TatA-depleted cells (Figure 2C). We observed a significant difference between the average AUC of normalized medial fluorescence profiles of WT and TatA-replete cells (Figure 2D), but this difference was small and may not be biologically relevant, as it may be due to ectopic and elevated expression of both TatA and mCherry-MreB. Similarly, there was no difference in mCherry-PBP1a accumulation in the midcell region upon osmotic upshift in WT, TatA-replete, and TatA-depleted cells (Figure S3, E and F). Together, these results indicate that, unlike PBP2, MreB and PBP1a do not depend on the Tat complex for subcellular localization, congruent with the expectation that, because MreB is a cytosolic protein (Figge *et al*., 2004) and PBP1a lacks a Tat signal sequence (Teufel *et al*., 2022), both are unlikely to interact directly with the Tat complex.

### TatA is essential for localization of PBP2 and ssTorA to the cell envelope

To verify that the Tat complex contributes to PBP2 subcellular localization even in the absence of osmotic shock, we imaged P*_van_-mCherry-pbp2* cells grown in PYE media as above and then immobilized on PYE agarose pads. We observed that mCherry-PBP2 localized predominantly to the cell periphery in WT and TatA-replete cells, consistent with insertion into the membrane; in contrast, TatA-depleted cells grown with glucose exhibited fluorescence throughout the cell body (Figure 3A). Plots of normalized fluorescence profiles along the transverse axis (across the cell width) of WT and TatA-replete cells displayed two peaks, corresponding to localization to the cell membrane on either side of the cell, whereas TatA-depleted cells showed a single central peak, suggesting cytosolic localization (Figure 3B). Measurements of normalized fluorescence at the midpoints of the transverse profiles indicated significantly higher levels in TatA-depleted cells compared to WT and TatA-replete cells (Figure 3B), confirming results from osmotic upshift experiments showing that PBP2 localization depends on TatA. We used antibodies against mCherry to examine steady-state levels of mCherry-PBP2 by Western blot analysis and detected bands corresponding to full-length mCherry-PBP2 as well as potentially cleaved forms (labeled as mCh-PBP2*), congruent with previous reports that *Caulobacter* PBP2 may have at least two forms (Divakaruni *et al*., 2005; Figge *et al*., 2004) (Figure 3C). Protein levels were relatively similar between cells replete with or depleted of TatA (or TatC) (Figure 3C, lanes 5-8); thus, the change in mCherry-PBP2 localization in Tat-depleted cells is likely due to improper membrane insertion, not variations in protein abundance.

**Figure 3.**
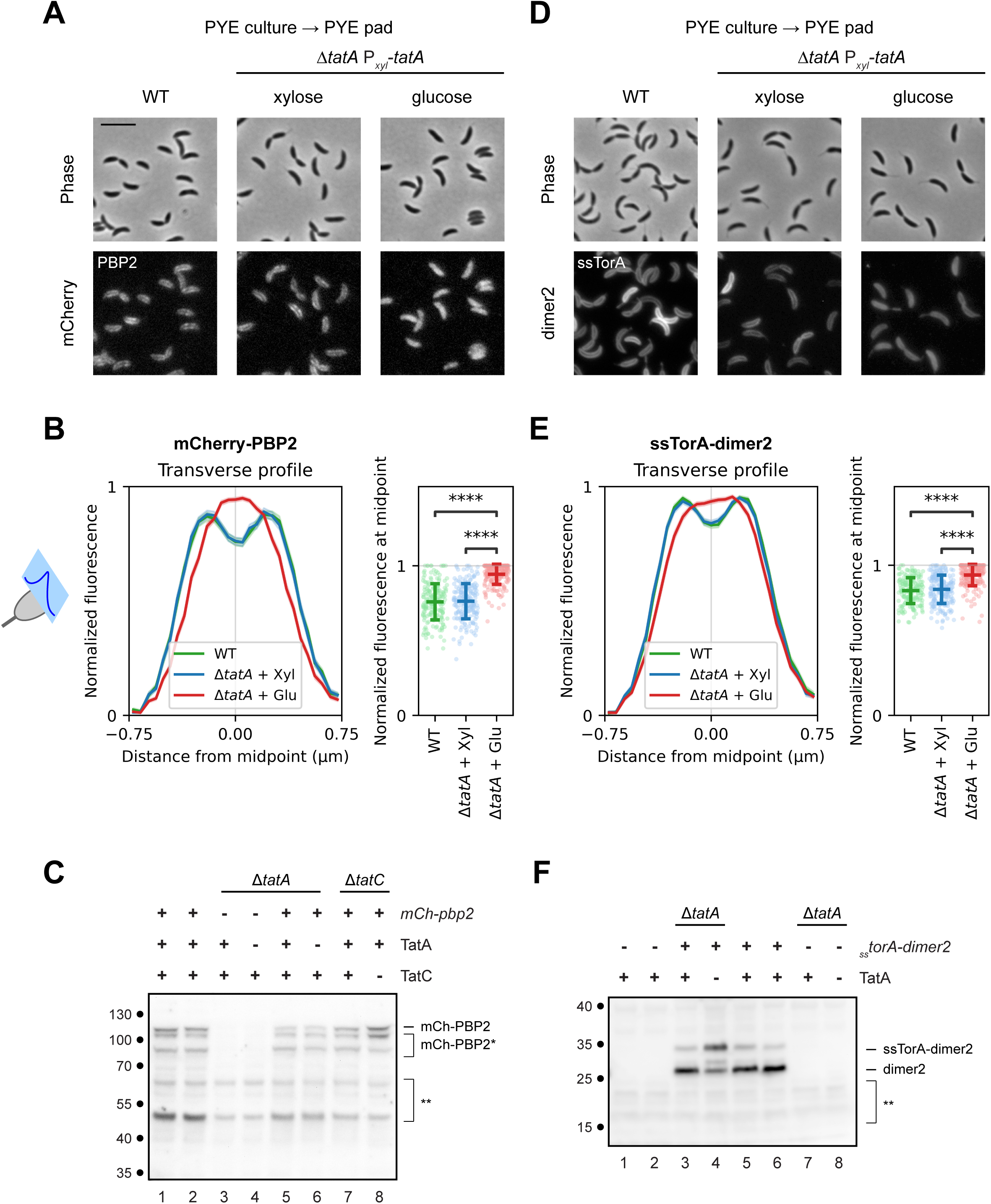
TatA is essential for targeting mCherry-PBP2 and ssTorA-dimer2 to the cell envelope. (**A**-**C**) WT and TatA depletion strains were induced with vanillate to express mCherry-PBP2, or (**D**-**F**) they constitutively express ssTorA-dimer2 from a plasmid. (**A**, **D**) Cells were grown in the presence or absence of glucose or xylose, as described in Figure 2, and transferred from PYE medium to PYE agarose pads for microscopy, without being subject to osmotic shock. Representative phase contrast (top) and fluorescence (bottom) images are shown. Scale bar, 5 µm. (**B**, **E**) Transverse profiles (left panels) represent normalized fluorescence intensities along the cell width (minor axis), as illustrated by the schematic of a model cell on the left. Colored lines indicate averages, while shaded areas indicate 95% confidence intervals. Normalized fluorescence at the midpoint of each transverse profile is shown at the population level as scatter plots (right panels), with horizontal bars indicating means and standard deviations. 400 cells were measured for each population; ****, *p* < 0.0001; based on two-tailed *t*-test. (**C**) Immunoblot probed with anti-mCherry antibodies to detect mCherry-PBP2. Cells were grown as above, with vanillate to induce expression of mCherry-PBP2 and xylose (odd-numbered lanes) or glucose (even-numbered lanes) to regulate expression of TatA or TatC. Labels above blot indicate relevant genotypes, presence (+) or absence (-) of *mCherry-pbp2*, and expression (+) or depletion (-) of TatA or TatC in the cells associated with each lane. Strains used were JOE3134 (WT background) (lanes 1, 2); JOE2723 (TatA depletion, without *mCherry-pbp2*) (lanes 3, 4); JOE3149 (TatA depletion) (lanes 5, 6); and JOE3147 (TatC depletion) (lanes 7, 8). All strains except for JOE2723 carried the P*_van_-mCherry-pbp2* allele. Positions of bands representing full-length mCherry-PBP2 (mCh-PBP2) and its derivatives (mCh-PBP2*), as well as non-specific bands (**), are indicated to the right of the blot. (**F**) Immunoblot probed with anti-mRFP1 antibodies to detect ssTorA-dimer2. Strains were grown with xylose (odd-numbered lanes) or glucose (even-numbered lanes) to regulate expression of TatA. Labels above blot indicate wild-type or Δ*tatA* background, presence (+) or absence (-) of the pEJ216 plasmid carrying *_ss_torA-dimer2*, and expression (+) or depletion (-) of TatA in the cells associated with each lane. Strains used were LS2677 (WT with vector) (lanes 1, 2); JOE7570 (TatA depletion with pEJ216) (lanes 3, 4); JOE7568 (WT with pEJ216) (lanes 5, 6); and JOE2723 (TatA depletion without plasmid) (lanes 7, 8). Positions of bands representing full-length ssTorA-dimer2 and its processed form (dimer2), as well as non-specific bands (**), are indicated to the right of the blot. Approximate molecular mass, in kDa, are shown to the left of the blots. Each blot is representative of two biological replicates.

By comparison, mCherry-MreB localized to the division site in the absence of osmotic upshift, as previously reported (Hocking *et al*., 2012), and this localization pattern did not depend on TatA (Figure S4A). Normalized fluorescence profiles along the transverse axis showed similar, single peaks for WT, TatA-replete, and TatA-depleted cells (Figure S4B). As for mCherry-PBP1a, we saw subtle localization to the cell periphery in some cells but mostly diffuse or patchy fluorescence throughout the cell body, regardless of TatA levels (Figure S4C). Transverse fluorescence profiles also all showed single broad peaks (Figure S4D). While PBP1a is a membrane protein (Strobel *et al*., 2014) and MreB associates with membrane proteins and the cell membrane (Alyahya *et al*., 2009; Divakaruni *et al*., 2007; Figge *et al*., 2004; White *et al*., 2010), we did not detect two prominent peaks in their transverse fluorescence profiles, likely due to low signal intensities, especially in regions away from the division site in the case of mCherry-MreB. In contrast to observations with PBP2, we did not see any significant difference in normalized fluorescence levels at the middle of these transverse profiles among WT, TatA-replete, or TatA-depleted cells (Figure S4, B and D), again suggesting that localization of MreB and PBP1a is independent of the Tat complex.

Finally, to confirm that TatA depletion in *Caulobacter* disrupts protein translocation by the Tat complex, we assessed localization of a fusion of the fluorescent protein dimer2 (Campbell *et al*., 2002) to the canonical Tat signal sequence of *E. coli* trimethylamine N-oxide reductase (ssTorA) (Cristóbal *et al*., 1999). This ssTorA-dimer2 fusion was previously shown to target to the periplasm when expressed from a plasmid in *Caulobacter* (Judd *et al*., 2005). As expected, WT and TatA-replete cells expressing the fusion exhibited fluorescence predominantly in their perimeters, while TatA-depleted cells displayed more diffuse fluorescence (Figure 3D), parallel to observations with mCherry-PBP2. Analysis of transverse fluorescence profiles (Figure 3E) also yielded similar results as mCherry-PBP2, suggesting a shift from periplasmic to cytoplasmic accumulation of ssTorA-dimer2 when TatA is depleted. Western blot analysis using antibodies against mRFP1 (a sibling of dimer2) revealed two bands in TatA-replete and WT cells, corresponding to full-length ssTorA-dimer2 and its processed form, dimer2, with the signal sequence removed following export into the periplasm (Figure 3F, lanes 3, 5, 6), as previously published (Judd *et al*., 2005). In TatA-depleted cells, the full-length form became more abundant, and an additional band appeared between those of the mature and processed forms, likely representing a degradation product in the cytoplasm (Figure 3F, lane 4). These results indicate that the *Caulobacter* Tat apparatus can recognize a known, heterologous Tat signal sequence for export. Moreover, similarities in the localization patterns of ssTorA-dimer2 and mCherry-PBP2 in the presence and absence of TatA suggest that PBP2 is a substrate of the Tat complex.

Together, our findings indicate that translocation and localization of the cell wall remodeling protein PBP2 depend on the essential Tat complex in *Caulobacter*. Only a few proteins that share the same topology as PBP2 (with a cytoplasmic N-terminus and single transmembrane helix) have been shown to involve the Tat pathway for membrane insertion (Bachmann *et al*., 2006; De Buck *et al*., 2007; James *et al*., 2013; Passmore *et al*., 2020). Our results potentially expand that repertoire. Furthermore, PBP2 is predicted to contain a Tat recognition sequence in a select number of model alpha-proteobacteria, including *Wolbachia pipientis*, *Rhodospirillum rubrum*, *Zymomonas mobilis*, and *Rhodobacter sphaeroides* (Figure 4 and Data Set S1), representing the orders *Rickettsiales*, *Rhodospirillales*, *Sphingomonadales*, and *Rhodobacterales*, respectively (Brilli *et al*., 2010; Meier-Kolthoff *et al*., 2022). A notable exception is the *Rhizobiales/Hyphomicrobiales* group, which lacks PBP2 and MreB (Brown *et al*., 2012; Cameron *et al*., 2014; Williams *et al*., 2021) and includes both *S. meliloti* and *A. tumefaciens*, in which the Tat system is essential and non-essential, respectively. We did not find Tat signal sequences in PBP2 orthologs of representative species outside of *Alphaproteobacteria* (Figure 4 and Data Set S1). Considering that the membrane anchor of PBP2 interfaces with RodA in *E. coli* (Liu *et al*., 2020; Nygaard *et al*., 2023; Shlosman *et al*., 2023), how the constraints of that interaction are reconciled with recognition by the Tat machinery in alpha-proteobacteria remains to be elucidated.

**Figure 4.**
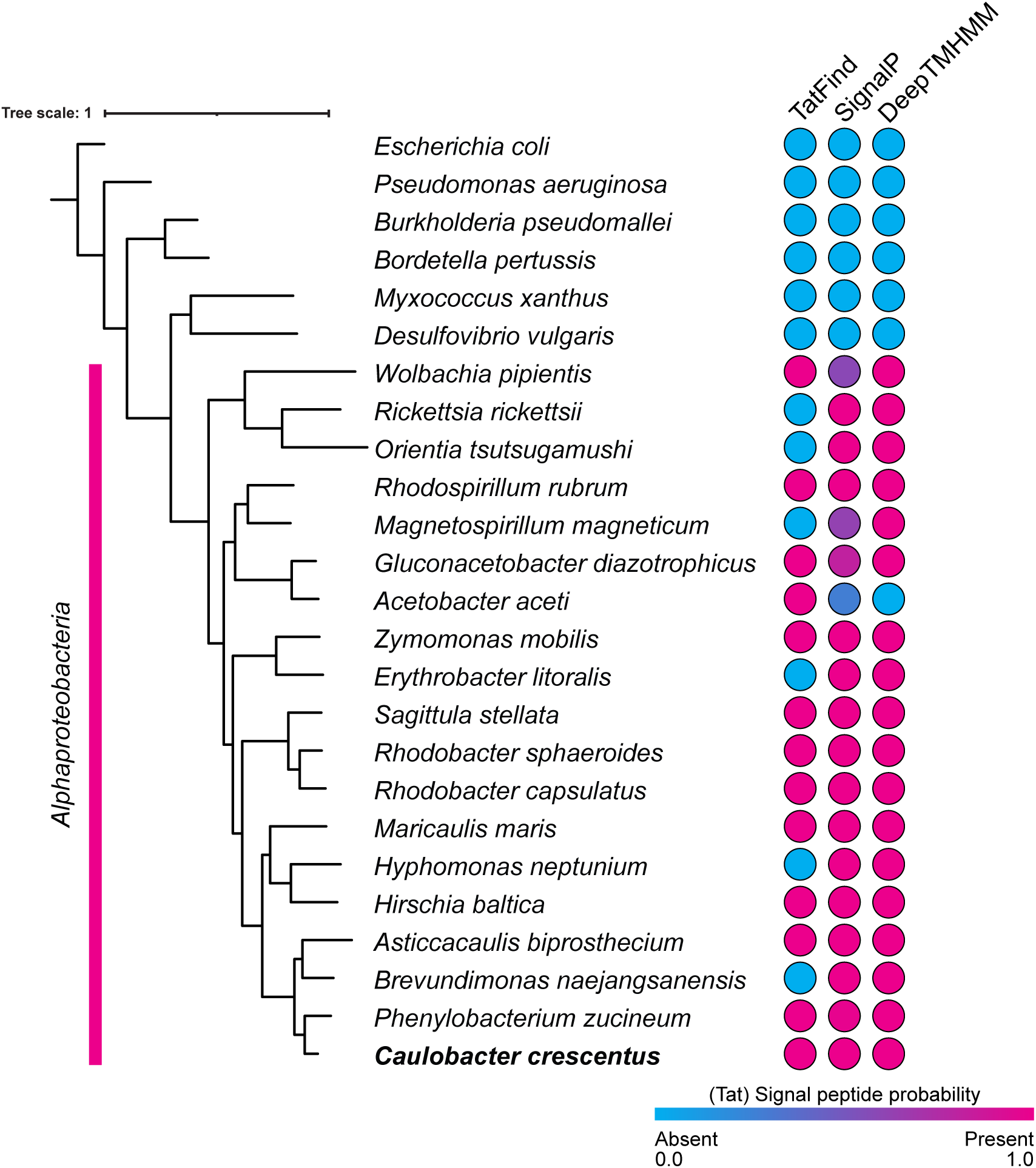
Conservation of Tat recognition sequences in PBP2 orthologs across *Alphaproteobacteria*. Three different signal sequence prediction tools (TatFind, SignalP, and DeepTMHMM) were used to determine the probability of Tat signals in PBP2 orthologs of select alpha-proteobacteria and other proteobacteria. All members of *Alphaproteobacteria* tested contain Tat recognition sequences within their PBP2, according to either TatFind or SignalP, and DeepTMHMM detected signal peptides in all except *Acetobacter aceti*. No representatives from *Rhizobiales* are shown because they lack *pbp2*. None of the algorithms found signal sequences in PBP2 outside of *Alphaproteobacteria*. Phylogenomic tree of shown species was constructed using GTDB-Tk.

While our study suggests that the Tat complex is necessary for PBP2 localization to the cell envelope in *Caulobacter*, further investigation is needed to determine if PBP2 is inserted into the membrane primarily via the Tat pathway. Although direct translocation is the simplest explanation, we cannot rule out that PBP2 translocation may involve cooperation between the Tat and Sec pathways (Keller *et al*., 2012; Tooke *et al*., 2017) or a hitchhiking mechanism with another Tat substrate (Gallego-Parrilla *et al*., 2024; Rodrigue *et al*., 1999). More work is also needed to determine if translocation by the Tat machinery is necessary for PBP2 to function in *Caulobacter*: for example, whether PBP2 folding needs to occur in the cytoplasm prior to export. Finally, although Tat-dependent localization of PBP2 may be a significant contributor to the essentiality of the Tat complex, many other proteins are predicted to be exported via the Tat pathway in *Caulobacter* (Dilks *et al*., 2003). Aside from PBP2, they are not known to participate directly in cell wall synthesis, but any number of them may be necessary for viability. A careful assessment of the essentiality of these potential substrates and their dependence on the Tat complex is needed to elucidate the critical nature of this export system. Understanding the role of the Tat pathway in *Caulobacter* may help reveal why select species have evolved to rely more than others on this system to secrete a significant number of proteins.

## Materials and Methods

### Bacterial strains, growth conditions, and molecular biology procedures

*C. crescentus* CB15, NA1000, and their derivatives were grown in PYE media with appropriate supplements, as published previously (Arellano *et al*., 2010; Chen *et al*., 2005). Standard protocols were used for strain and plasmid construction, including cloning, manipulation, and analysis of DNA, also as previously described (Chen *et al*., 2005; Fields *et al*., 2012). Detailed descriptions of strains and plasmids, their construction, and primer sequences are provided in the supplemental materials.

### Growth assays

For growth curves in liquid media, stationary overnight cultures in PYE with 0.1% xylose were washed twice with plain PYE and diluted to an OD600 of 0.01 in PYE with either 0.1% xylose or 0.1% glucose. 150 µL of diluted cell culture was added to Corning transparent 96-well plates, which were covered with Breathe-Easy membranes to prevent excessive evaporation. A Tecan Genios plate reader was used to measure OD610 every 15 minutes over a period of 24 hours with shaking in between readings at 30°C. MATLAB was used to plot the average OD610 across two independent experiments, each with four replicate wells (8 wells total), with the standard deviation denoted as the shaded area. For visualization of cell morphology, cultures were grown similarly in a BioTek Synergy H1 plate reader in 96-well plates with plastic lids (instead of membranes). Colonies on agar plates were photographed with digital cameras.

### Microscopy

For visualization of cell morphology, aliquots (2 – 3 µL) of cultures grown in 96-well plates (see above) were spotted on 1% agarose pads made with PYE. Bacterial cells were examined by phase contrast microscopy using a Zeiss Axio Imager M1 with a 63x Plan-Apochromat objective (numerical aperture of 1.4), and images were acquired with a Zeiss AxioCam MRm camera and AxioVision software, as previously described (Fields *et al*., 2012).

For fluorescent protein localization, WT or Tat depletion strains were grown overnight in PYE with 0.1% xylose. Overnight cultures were washed twice with plain PYE and diluted to an OD600 of 0.1 in PYE with 0.5 mM vanillate and either 0.1% xylose or 0.1% glucose and grown for four hours. For depletion of TatC, strains were diluted to an OD600 of 0.1 and grown to an OD600 of 0.8 in PYE with 0.1% xylose, washed twice with plain PYE, and diluted to an OD600 of 0.002 - 0.004 in PYE with 0.1% glucose. After 14 - 16 hours of growth, the cultures were diluted to an OD600 of 0.1 in PYE with 0.5 mM vanillate and 0.1% xylose or glucose and grown for another four hours. Approximately 2 µL of cell culture was then spotted on 1% agarose pads made with either PYE (Poindexter, 1964) or M2 (Johnson & Ely, 1977). Micrographs were captured using the same Zeiss Axio Imager M1 system described above, with an HQ Texas Red filter set 45 (excitation band pass, 560/40 nm; beam splitter, 585 nm; emission band pass, 630/75 nm). Typical exposure for fluorescence images was 1 s. Each experiment was repeated at least twice.

### Image Analysis

For analysis of cell morphology, MicrobeJ (Ducret *et al*., 2016) was used to identify bacterial cells in the phase contrast images. After the outputs were manually curated to eliminate errors (such as lysed cells, debris, or two cells grouped as one), the “SHAPE.roundness” values were obtained.

For fluorescence measurements along the medial axis (along the length, or major axis, of each cell), MicrobeJ was again used to identify bacterial cells in the phase contrast images and record the medial profiles from the fluorescence channel. After the outputs were manually curated to eliminate errors, demographs were generated using the normalized medial profiles. The normalized medial profiles were interpolated to 21 points to obtain the relative fluorescence along the normalized cell length. Area under curve for each medial profile was estimated using the trapezoidal rule, by calculating the areas of 20 trapezoids fitted under the plot.

For measurements along the transverse axis (the minor axis), a line was drawn manually in ImageJ (Schneider *et al*., 2012) across the widest section of each cell, with each end of the line 0.75 μm from the midpoint of the cell. The fluorescence intensity along the line was normalized, and normalized fluorescence intensities at midpoint were then compared for statistical significance.

All plots were generated using matplotlib/seaborn (Hunter, 2007; Waskom, 2021), and *p*-values were calculated using SciPy two-tailed independent *t*-test (Virtanen *et al*., 2020).

### Western blotting

Immunoblotting was performed essentially as previously described (Chen *et al*., 2005; Judd *et al*., 2005). Cells from 2-mL aliquots of cultures were harvested and resuspended in SDS sample buffer (200 µL for OD600 of 1) and boiled. 10 µL of each sample was used for SDS-PAGE, and proteins were transferred onto PVDF membrane. Immunodetection was done with rabbit polyclonal antibodies against mCherry (Invitrogen PA5-34974) or mRFP1 (Chen *et al*., 2005), peroxidase-conjugated donkey anti-rabbit secondary antibodies (Thermo 31460), and chemiluminescent reagents (SuperSignal West Pico).

### Sequence analysis

PBP2 orthologs were identified by BLAST in UniProt using PBP2 from *C. crescentus* as the query and, when necessary, verified by annotation or proximity of the corresponding gene to *rodA* (UniProt, 2025). Protein sequences were analyzed using <u>TatFind</u>, <u>SignalP 6.0</u>, and <u>DeepTMHMM 1.0</u> (Dilks *et al*., 2003; Hallgren *et al*., 2022; Rose *et al*., 2002; Teufel *et al*., 2022) (Figure 4, Data Set S1). TatFind and DeepTMHMM produced binary outputs for the detection of Tat signal sequence and any signal sequence or transmembrane segment, respectively. All PBP2 orthologs chosen from species outside of *Alphaproteobacteria* were predicted by DeepTMHMM to have transmembrane segments instead of signal peptides. SignalP provided probabilities for Tat/SPI, and Tat/SPII (lipoprotein); for the purpose of this analysis, these values were combined to generate a total Tat probability. Alignments of signal peptides was performed using MAFFT and Jalview (Katoh & Standley, 2013; Waterhouse *et al*., 2009), and amino acid color is based on the hydrophobicity of a given residue (Data Set S1). Whole-genome sequences for the sampled species were acquired from GenBank (Sayers *et al*., 2025), and phylogenomic tree was generated using GTDB-Tk’s identify, align, and infer tools with default parameters; branch length is based on relative evolutionary divergence and average nucleotide identity, under the Whelan-Goldman (WAG) model (Chaumeil *et al*., 2022).

## Supporting information

Data Set S1

## Abbreviations used

Tat: twin-arginine transport
PBP: penicillin-binding protein
PG: peptidoglycan
OD: optical density
AUC: area under the curve

## Acknowledgements

We would like to thank Lucy Shapiro for her encouragement and generosity, as well as past members of the Lucy Shapiro, Harley McAdams, and Joseph Chen labs for their help in this investigation. K.C., N.P., and D.L.W. received support from the Genentech Foundation Scholars Program. In addition, D.L.W. was supported by the NIH MS Bridges to the Doctorate Program under Award Number R25-GM048972, and N.P. was supported by the NIH U-RISE Program under Award Number T34-GM145400. J.C.C. has received support from the NIGMS of the NIH under award numbers F32 GM067472, SC2 GM082318, and SC3 GM096943. We thank two *Molecular Biology of the Cell* reviewers for their insightful feedback on how to improve the manuscript. The content is solely the responsibility of the authors and does not necessarily represent the official views of the funding agencies.

