## Supplemental Figures S1, S2, S3, S4 for "Twin-arginine transport complex plays an essential role in *Caulobacter* cell shape and viability"

### Figure S1.

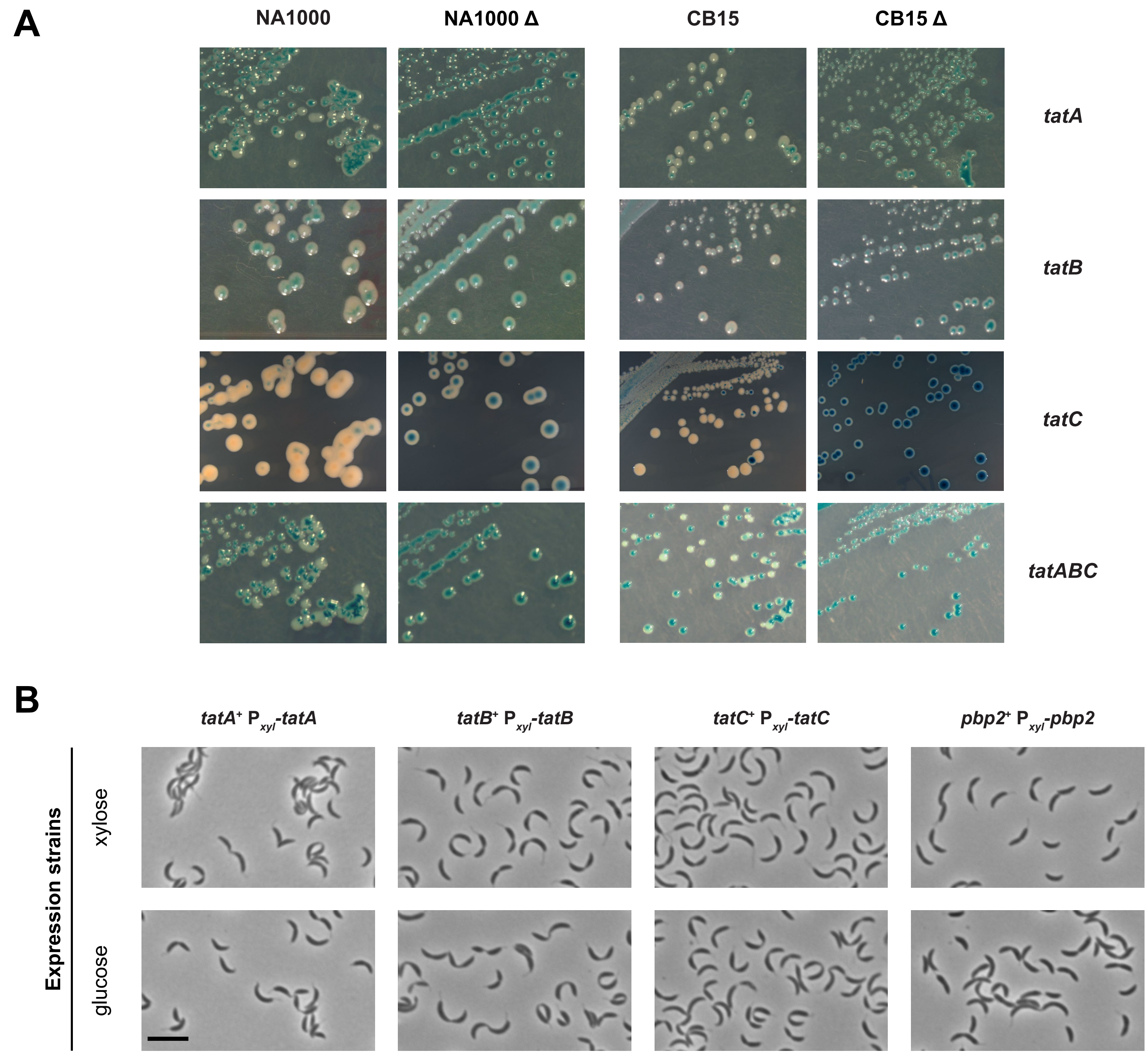


**Figure S1.** Colonies and cell morphology of *tat* and *pbp2* strains. **(A)** Plasmid loss and colony-sectoring assay. Strains with deletions of *tat* genes are unable to lose their complementing plasmids. NA1000- or CB15-derived strains were constructed with wild-type alleles or deletions of *tat* genes (Δ) on the chromosome while carrying the corresponding *tat* genes and the *E. coli lacZ* gene on complementing plasmids. They were streaked onto PYE plates containing X-Gal and incubated at 30°C. Resultant colonies were photographed, sometimes after plates were refrigerated for several days to enhance the blue color. Images shown here demonstrate the variations in colony and background colors due to factors such as incubation period and camera lighting, but all indicate that deletion strains do not produce white colonies because they are unable to lose the *lacZ* gene, linked to the complementing *tat* allele. Strains used were constructed from NA1000 [JOE2670 and JOE2692 (*tatA*), JOE2713 and JOE2712 (*tatB*), JOE2402 and JOE2406 (*tatC*), JOE2856 and JOE2877 (*tatABC*)] or CB15 [JOE2674 and JOE2696 (*tatA*), JOE2749 and JOE2748 (*tatB*), JOE2403 and JOE2408 (*tatC*), JOE2858 and JOE2890 (*tatABC*)]. Images are representative of two biological replicates. **(B)** Cell morphology of wild-type strains expressing *tat* or *pbp2* from the *xylX* promoter ($P_{xyl}$). Phase contrast images were acquired after expression strains were grown under inducing (xylose) or non-inducing (glucose) conditions for 22-24 hours (*n* = 3 biological replicates). Strains used were JOE2719, JOE2758, JOE2332, and JOE7294. Scale bar, 5 µm.

### Figure S2.

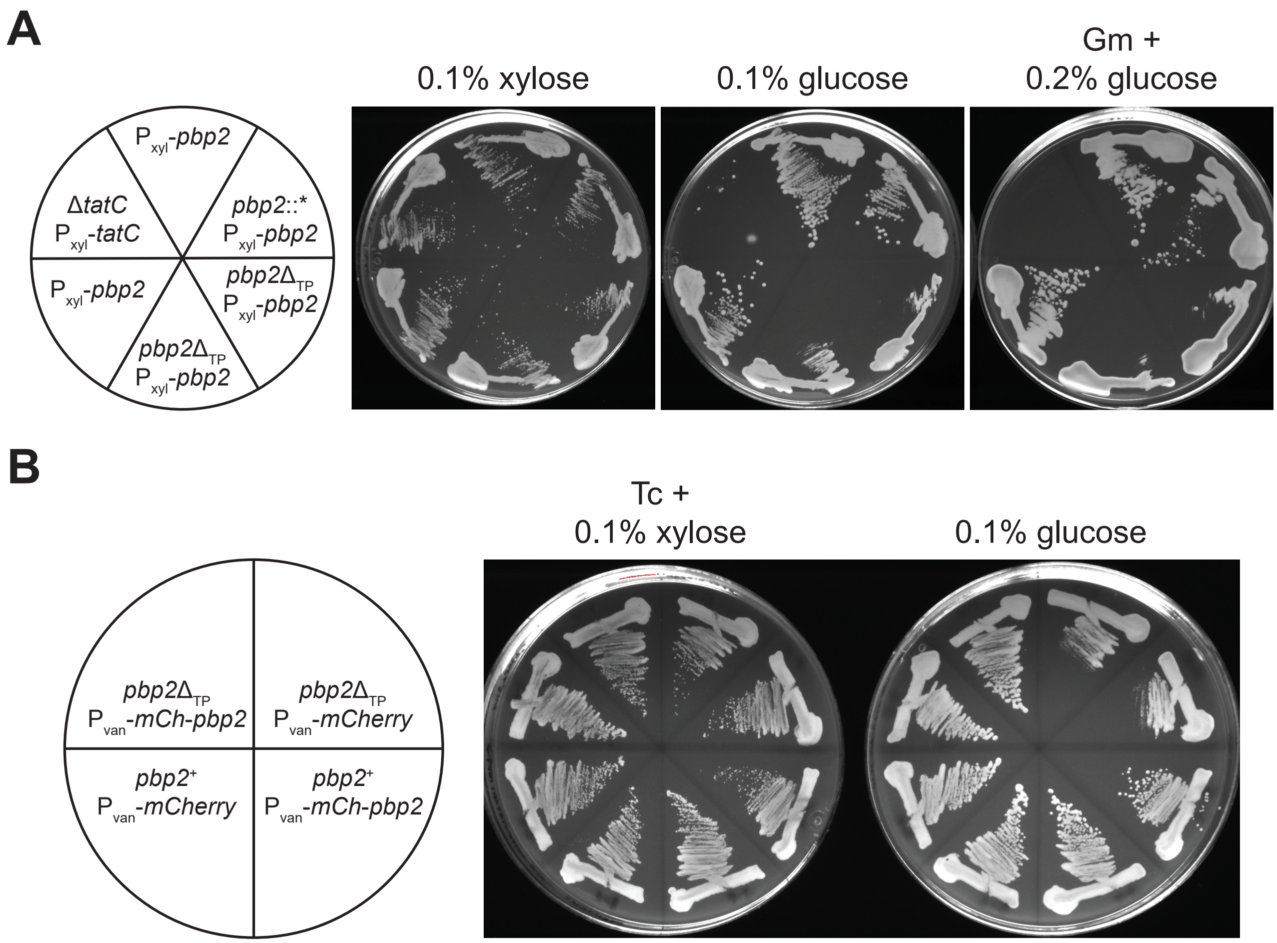


**Figure S2.** PBP2 is essential for colony formation in *Caulobacter*. (**A**) PBP2 is required for robust colony formation. Strains were streaked onto PYE plates containing 0.1% xylose, 0.1% glucose, or 5 µg/mL gentamicin (Gm) + 0.2% glucose. Relevant genotypes of strains in each sector are shown in the schematic on the left. Strains used were JOE2357, JOE7295, JOE7296, JOE7301, JOE7302, and JOE7303. JOE7296 (*pbp2*::* $P_{xyl}$-*pbp2*) is derived from JOE7295 and carries the allelic replacement plasmid pTC358 integrated into the *pbp2* locus. Strains with deletion (JOE7301 and JOE7302) or wild-type (JOE7303) allele of *pbp2* were derived from JOE7296 following sucrose counter-selection. Images represent results from three or more independent replicates, each done on a separate day. (**B**) *mCherry-pbp2* can complement null mutation in *pbp2*. Strains with or without the *pbp2*$\Delta_{TP}$ mutation and expressing mCherry or mCherry-PBP2 from the vanillate-inducible promoter ($P_{van}$) were streaked onto PYE plates containing 0.1% xylose + 1 µg/mL oxytetracycline (Tc) or 0.1% glucose to induce or repress, respectively, $P_{xyl}$-*pbp2*. Basal expression of mCherry-PBP2 from the vanillate promoter is sufficient for complementation. Schematic on the left indicate relevant genotypes of strains in each quadrant. Strains used were JOE7324, JOE7326, JOE7328, and JOE7329. Image shown represents results from three or more independent trials.

### Figure S3.

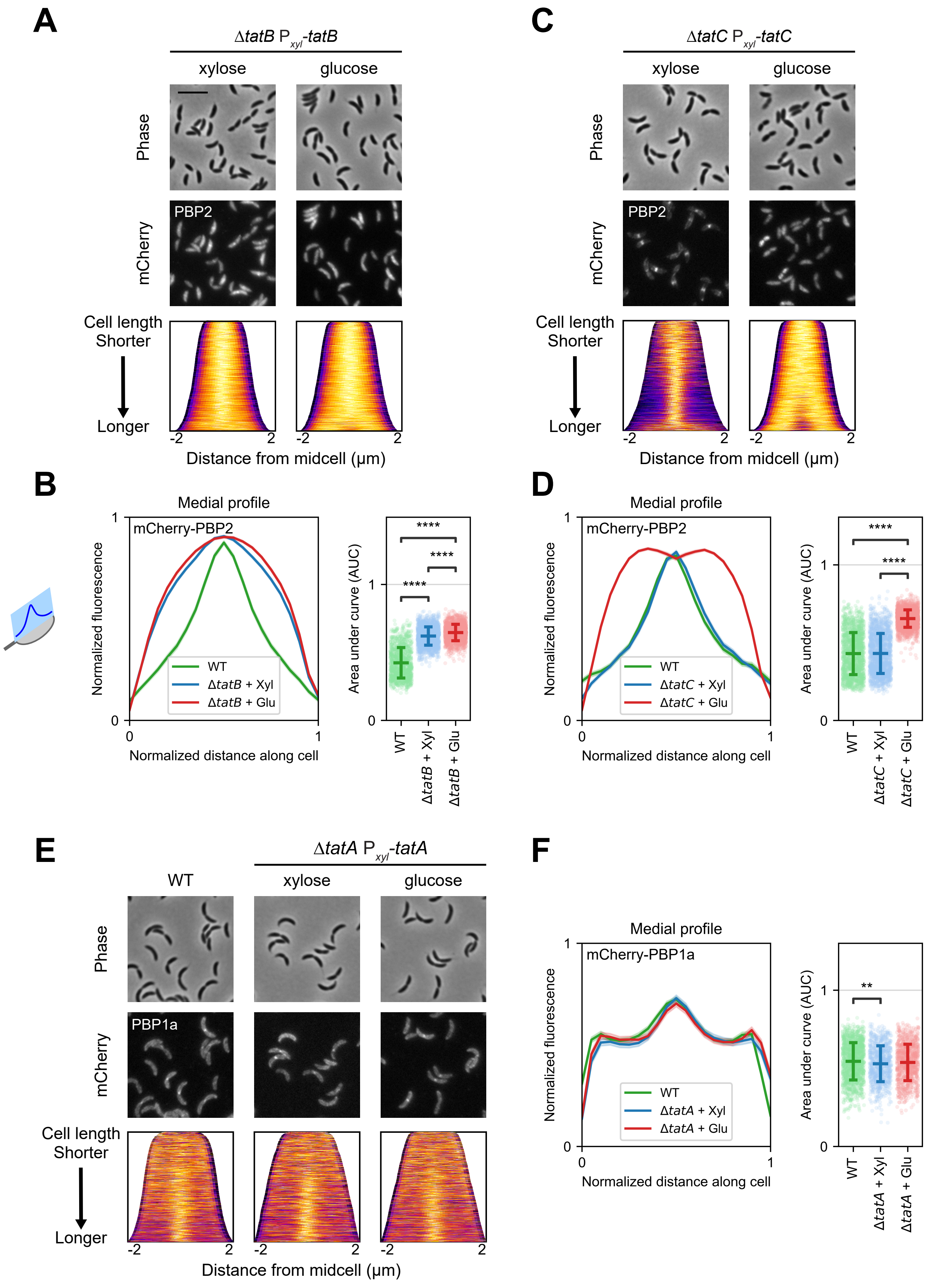


**Figure S3.** Midcell localization of PBP2 and PBP1a in Tat depletion strains upon osmotic shock upshift.

### Figure S3.

**Figure S3.** Midcell localization of PBP2 and PBP1a in Tat depletion strains upon osmotic shock upshift. Cultures were induced for four hours with vanillate to express (**A**, **B**, **C**, **D**) mCherry-PBP2 or (**E**, **F**) mCherry-PBP1a. (**A**, **B**) TatB depletion strain (Δ*tatB* $P_{xyl}$-*tatB*) was grown in the presence of xylose or glucose for four hours to express or repress *tatB*, respectively, while (**C**, **D**) TatC depletion strain (Δ*tatC* $P_{xyl}$-*tatC*) was first grown for 14 – 16 hours with glucose and then resuspended in PYE medium containing xylose or glucose for four additional hours of growth. (**E**, **F**) WT and TatA depletion strains expressing mCherry-PBP1a were grown similarly as those in Figure 2, in the absence or presence of xylose or glucose for four hours. Cells were subjected to osmotic upshift from being transferred from PYE medium to an M2 agarose pad for microscopy. (**A**, **C**, **E**) Representative phase contrast (top) and fluorescence (middle) images are shown with corresponding population-level demographs (bottom). Demographs depict localization of normalized fluorescence along the medial axis (the cell length), with cells arranged by length and lighter colors indicating brighter fluorescence. (**B**, **D**, **F**) Medial profiles (left panels) represent normalized fluorescence intensities along normalized cell length, as illustrated by the schematic of a model cell to the left of part (B). Colored lines indicate averages, while shaded areas indicate 95% confidence intervals. Area under curve was calculated for each medial profile and shown at the population level as scatter plots (right panels), with horizontal bars indicating means and standard deviations. (**A**, **B**) Midcell localization of mCherry-PBP2 is ineffective in the TatB depletion strain, regardless of TatB expression; n = 1146 (WT), 1853 (Δ*tatB* + Xyl), 1611 (Δ*tatB* + Glu). (**C**, **D**) Midcell localization of mCherry-PBP2 depends on TatC; n = 1568 (WT), 1683 (Δ*tatC* + Xyl), 2126 (Δ*tatC* + Glu). (**E**, **F**) Midcell localization of mCherry-PBP1a is not affected by TatA depletion; n = 1415 (WT), 877 (Δ*tatA* + Xyl), 991 (Δ*tatA* + Glu). Scale bar, 5 µm. **, $p < 0.01$; ****, $p < 0.0001$; based on two-tailed *t*-test.

### Figure S4.

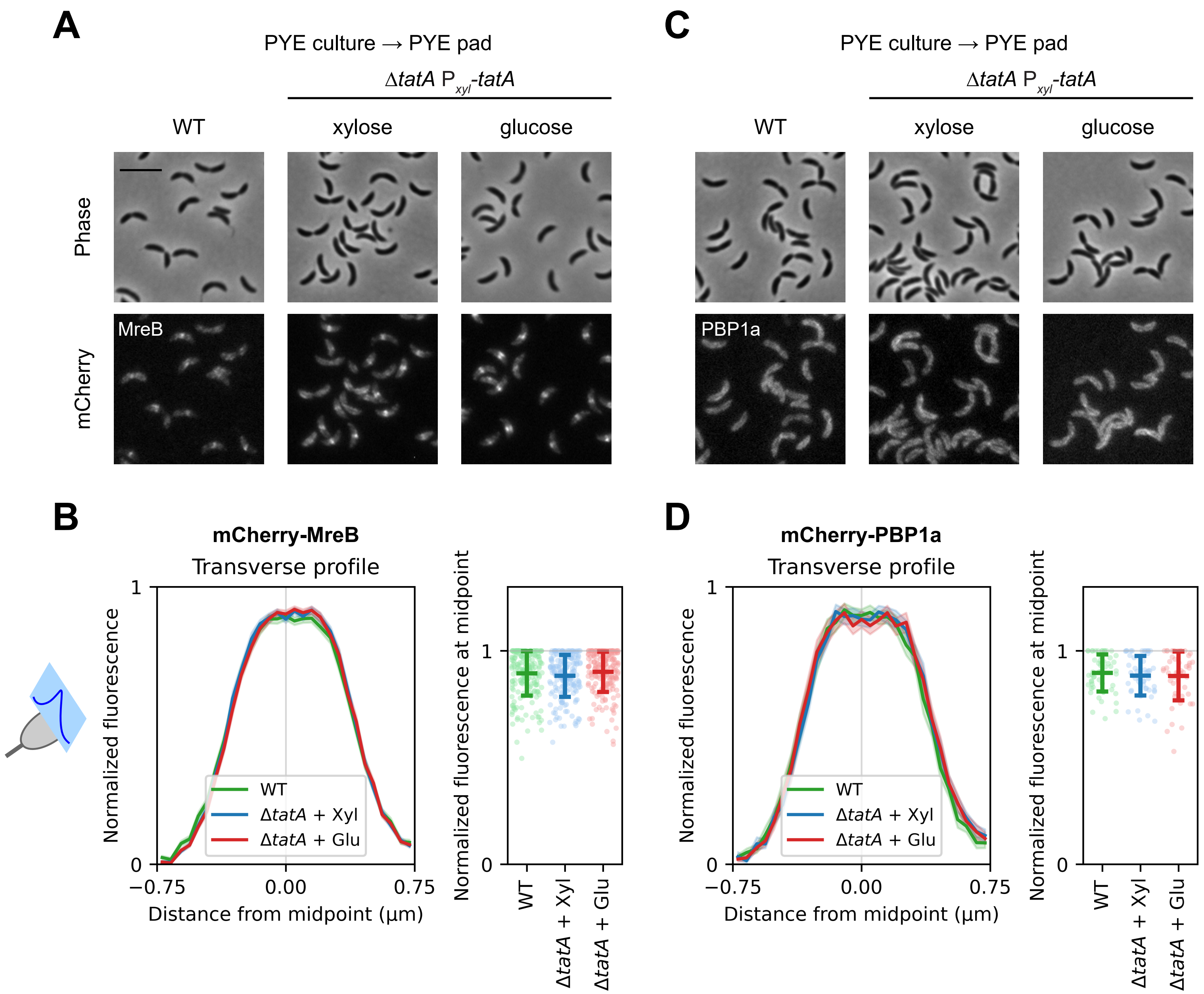


**Figure S4.** Localization of MreB and PBP1a without osmotic upshift. WT and TatA depletion strains were induced with vanillate to express (**A**, **B**) mCherry-MreB or (**C**, **D**) mCherry-PBP1a and grown in the absence or presence of xylose or glucose, as described in Figure 2, and transferred from PYE medium to PYE agarose pads for microscopy, without being subject to osmotic shock. (**A**, **C**) Representative phase contrast (top) and fluorescence (bottom) images are shown. Scale bar, 5 μm. (**B**, **D**) Transverse profiles (left panels) represent normalized fluorescence intensities along the cell width (minor axis), as illustrated by the schematic of a model cell on the left. Colored lines indicate averages, while shaded areas indicate 95% confidence intervals. Normalized fluorescence at the midpoint of each transverse profile is shown at the population level as scatter plots (right panels), with horizontal bars indicating means and standard deviations. 400 cells were measured for each mCherry-MreB population, while 100 cells were measured for each mCherry-PBP1a population. No significant differences were observed among populations with the same fluorescence fusion, based on two-tailed *t*-tests.
