## Supplementary material for "Twin-arginine transport complex plays an essential role in *Caulobacter* cell shape and viability": Tables S1-S3

**Supplemental Materials**

List of contents:

**Table S1.** Strains used for this study of Tat system.
**Table S2.** Plasmids used for this study.
**Table S3.** Primers used for this study.

**Supplemental Figure Legends**
Figure S1. Colonies and cell morphology of *tat* and *pbp2* strains.
Figure S2. PBP2 is essential for colony formation in *Caulobacter*.
Figure S3. Midcell localization of PBP2 and PBP1a in Tat depletion strains upon osmotic upshift.
Figure S4. Localization of MreB and PBP1a without osmotic upshift.

**Supplemental References**

**Data Set S1.** PBP2 orthologs in select proteobacteria.

Brief description:

Strains used and how they were generated are described in Table S1. Plasmids used and their constructions are provided in Table S2. Table S3 lists primer sequences. Figure legends for Supplemental Figures S1 – S4 are included as well, followed by references cited in the supplemental materials.

Data Set S1 contains sequences of PBP2 orthologs from select proteobacteria, along with predictions of Tat recognition sequences and signal peptides generated by TatFind, SignalP, and DeepTMHMM. The file also includes a multiple sequence alignment of the N-termini of the PBP2 orthologs, with residues color-coded by hydrophobicity, as well as conservation and consensus annotations.

**TABLE S1.** Strains used for this study of Tat system.

| Strains | Relevant genetic markers, features, and/or description [1] | Source / Notes [2] |
|---|---|---|
| CB15 | wild-type *C. crescentus* | (Poindexter, 1964) |
| NA1000 | *syn-1000*; previously called CB15N, a synchronizable derivative of CB15 | (Evinger & Agabian, 1977) |
| LS2677 | NA1000 / pJS14 ($Cm^R$) | (Judd *et al.*, 2005) |
| JOE2321 | NA1000 Δ*lacA* | (Arellano *et al.*, 2010) |
| JOE2323 | CB15 Δ*lacA* | (Arellano *et al.*, 2010) |
| JOE2332 | NA1000 *xylX*::pJC351 ($P_{xyl}$-*CC2001*; $Sp^R$) | mate pJC351 into NA1000 |
| JOE2402 | NA1000 Δ*lacA* / pJC363 ($P_{xyl}$-*CC2001*; $Cm^R$) | electroporate pJC363 into JOE2321 |
| JOE2403 | CB15 Δ*lacA* / pJC363 ($P_{xyl}$-*CC2001*; $Cm^R$) | electroporate pJC363 into JOE2323 |
| JOE2670 | NA1000 Δ*lacA* / pJC397 (*CC2003*; $Cm^R$) | electroporate pJC397 into JOE2321 |
| JOE2674 | CB15 Δ*lacA* / pJC397 (*CC2003*; $Cm^R$) | electroporate pJC397 into JOE2323 |
| JOE2713 | NA1000 Δ*lacA* / pJC396 ($P_{xyl}$-*CC2002*; $Cm^R$) | allelic replacement with pJC368 [3] |
| JOE2719 | NA1000 *xylX*::pJC399 ($P_{xyl}$-*CC2003*; $Sp^R$) | mate pJC399 into NA100 |
| JOE2749 | CB15 Δ*lacA* / pJC396 ($P_{xyl}$-*CC2002*; $Cm^R$) | allelic replacement with pJC368 [3] |
| JOE2758 | NA1000 Δ*lacA xylX*::pJC366 ($P_{xyl}$-*CC2002*; $Sp^R$) | mate pJC366 into JOE2321 |
| JOE2856 | NA1000 Δ*lacA* / pJC378 (*CC2003-CC2001*; $Tc^R$) | mate pJC378 into JOE2321 |
| JOE2858 | CB15 Δ*lacA* / pJC378 (*CC2003-CC2001*; $Tc^R$) | mate pJC378 into JOE2323 |
| JOE7294, JOE7295 | NA1000 *xylX*::pJC774 ($P_{xyl}$-*CC1546*; $Gm^R$) | electroporate pJC774 into NA1000 |
| **Sectoring (Plasmid Loss Detection) strains** | | |
| CC2001 (TatC) | | |
| JOE2406 | NA1000 Δ*lacA* Δ*CC2001*::Ω*aacC4* / pJC363 ($Cm^R$) | JOE2402xΦCr30(JOE2361), select for $Am^R$ |
| JOE2408 | CB15 Δ*lacA* Δ*CC2001*::Ω*aacC4* / pJC363 ($Cm^R$) | JOE2403xΦCr30(JOE2361), select for $Am^R$ |
| CC2002 (TatB) | | |
| JOE2712 | NA1000 Δ*lacA* Δ*CC2002* / pJC396 ($Cm^R$) | allelic replacement with pJC368 [3] |
| JOE2748 | CB15 Δ*lacA* Δ*CC2002* / pJC396 ($Cm^R$) | allelic replacement with pJC368 [3] |
| CC2003 (TatA) | | |
| JOE2647 | NA1000 Δ*lacA* Δ*CC2003*::*aacC4* / pJC392 ($Tc^R$) | allelic replacement with pJC372 [4] |
| JOE2692 | NA1000 Δ*lacA* Δ*CC2003*::*aacC4* / pJC397 ($Cm^R$) | JOE2670xΦCr30(JOE2647), select for $Am^R$ |
| JOE2696 | CB15 Δ*lacA* Δ*CC2003*::*aacC4* / pJC397 ($Cm^R$) | JOE2674xΦCr30(JOE2647), select for $Am^R$ |
| CC2003-CC2001 (TatA-TatC) | | |
| JOE2843 | NA1000 Δ*CC2003-CC2001*::*aacC4* / pJC378 ($Tc^R$) | allelic replacement with pJC402 [5] |
| JOE2877 | NA1000 Δ*lacA* Δ*CC2003-CC2001*::*aacC4* / pJC378 ($Tc^R$) | JOE2856xΦCr30(JOE2843), select for $Am^R$ |
| JOE2890 | CB15 Δ*lacA* Δ*CC2003-CC2001*::*aacC4* / pJC378 ($Tc^R$) | JOE2858xΦCr30(JOE2843), select for $Am^R$ |

| | | |
|---|---|---|
| **Depletion strains** | | |
| JOE2357 | NA1000 Δ*tatC xylX*::pJC351 ($P_{xyl}$-*CC2001*; Sp$^R$) | allelic replacement with pJC305 [6] |
| JOE2361 | NA1000 Δ*tatC*::Ω*aacC4 xylX*::pJC351 ($P_{xyl}$-*CC2001*; Sp$^R$) | allelic replacement with pJC349 [6] |
| JOE2475 | NA1000 Δ*tatB xylX*::pJC366 ($P_{xyl}$-*CC2002*; Sp$^R$) | allelic replacement with pJC368 [7] |
| JOE2723 | NA1000 Δ*tatA*::*aacC4 xylX*::pJC399 ($P_{xyl}$-*CC2003*; Sp$^R$) | JOE2719xΦCr30(JOE2647), select for Am$^R$ |
| JOE7298, JOE7302 | NA1000 *pbp2*$\Delta_{TP}$ *xylX*::pJC774 ($P_{xyl}$-*CC1546*; Gm$^R$) | allelic replacement with pTC358 [8] |
| JOE7303 | NA1000 *pbp2*$^+$ *xylX*::pJC774 ($P_{xyl}$-*CC1546*; Gm$^R$) | allelic replacement with pTC358 [8] |
| JOE7324 | NA1000 *pbp2*$\Delta_{TP}$ *xylX*::pJC774 *vanA*::pJC427 ($P_{van}$-*mCherry-pbp2*; Tc$^R$) | mate pJC427 into JOE7302 |
| JOE7326 | NA1000 *pbp2*$\Delta_{TP}$ *xylX*::pJC774 *vanA*::pVCHYN-5 ($P_{van}$-*mCherry*; Tc$^R$) | mate pVCHYN-5 into JOE7302 |
| JOE7328 | NA1000 *xylX*::pJC774 *vanA*::pJC427 ($P_{van}$-*mCherry-pbp2*; Tc$^R$) | mate pJC427 into JOE7303 |
| JOE7329 | NA1000 *xylX*::pJC774 *vanA*::pVCHYN-5 ($P_{van}$-*mCherry*; Tc$^R$) | mate pVCHYN-5 into JOE7303 |
| **Localization of MreB, PBP1a, PBP2, ssTorA-dimer2** | | |
| JOE3130 | NA1000 *vanA*::pJC424 ($P_{van}$-*mCherry-mreB*; Tc$^R$) | mate pJC424 into NA1000 |
| JOE3134 | NA1000 *vanA*::pJC427 ($P_{van}$-*mCherry-pbp2*; Tc$^R$) | mate pJC427 into NA1000 |
| JOE3143 | NA1000 Δ*tatA*::*aacC4 xylX*::pJC399 ($P_{xyl}$-*CC2003*; Sp$^R$) *vanA*::pJC424 ($P_{van}$-*mCherry-mreB*) | mate pJC424 into JOE2723 |
| JOE3147 | NA1000 Δ*tatC*::Ω*aacC4 xylX*::pJC351 ($P_{xyl}$-*CC2001*; Sp$^R$) *vanA*::pJC427 ($P_{van}$-*mCherry-pbp2*) | mate pJC427 into JOE2361 |
| JOE3148 | NA1000 Δ*tatB xylX*::pJC366 ($P_{xyl}$-*CC2002*; Sp$^R$) *vanA*::pJC427 ($P_{van}$-*mCherry-pbp2*) | mate pJC427 into JOE2475 |
| JOE3149 | NA1000 Δ*tatA*::*aacC4 xylX*::pJC399 ($P_{xyl}$-*CC2003*; Sp$^R$) *vanA*::pJC427 ($P_{van}$-*mCherry-pbp2*) | mate pJC427 into JOE2723 |
| JOE7559 | NA1000 *vanA*::pTC343 ($P_{van}$-*mCherry-pbp1a*; Tc$^R$) | mate pTC343 into NA1000 |
| JOE7562 | NA1000 Δ*tatA*::*aacC4 xylX*::pJC399 ($P_{xyl}$-*CC2003*; Sp$^R$) *vanA*::pTC343 ($P_{van}$-*mCherry-pbp1a*; Tc$^R$) | mate pTC343 into JOE2723 |
| JOE7568 | NA1000 / pEJ216 ($_{ss}$*torA-dimer2*; Cm$^R$) | electroporate pEJ216 into NA1000 |
| JOE7570 | NA1000 Δ*tatA*::*aacC4 xylX*::pJC399 ($P_{xyl}$-*CC2003*; Sp$^R$) / pEJ216 ($_{ss}$*torA-dimer2*; Cm$^R$) | electroporate pEJ216 into JOE2723 |

[1] Km, Tc, Sp, Cm, Am: abbreviations for kanamycin, (oxy)tetracycline, spectinomycin, chloramphenicol, apramycin.

[2] Φ indicates generalized transduction, as mediated by bacteriophage ΦCr30. For example, JOE2373 x Φ(JOE2361) means that a bacteriophage lysate made from JOE2361 was used to infect JOE2373.

[3] To obtain JOE2712, JOE2713, JOE2748, and JOE2749, pJC368 was mated into JOE2321 or JOE2323 for integration at the *CC2002* locus, and subsequently pJC396 was introduced by electroporation. The resulting strains were counter-selected with sucrose in the presence of xylose, and sucrose-insensitive isolates were screened for the presence or deletion of the native *tatB* gene.

[4] To obtain JOE2647, pJC372 was mated into JOE2321 for integration at the *CC2003* locus, followed by introduction of the pJC392 complementing plasmid, also via mating. The resulting strain was counter-selected with sucrose, and sucrose-insensitive isolates were screened for the replacement of the native *tatC* gene with *aacC4*, which confers resistance to apramycin.

[5] To obtain JOE2843, pJC402 was mated into NA1000 for integration into the chromosome, followed by introduction of the pJC378 complementing plasmid, also via mating. The resulting strain was counter-selected with sucrose, and sucrose-insensitive isolates were screened for the replacement of the native *tatABC* genes with *aacC4*, which confers resistance to apramycin.

[6] To obtain JOE2357 and JOE2361, pJC305 or pJC349 was mated into NA1000 for integration at the *CC2001* locus, and subsequently pJC351 was integrated at the *xylX* locus, also introduced by mating. The resulting strains were counter-selected with sucrose in the presence of xylose, and sucrose-insensitive isolates were screened for the deletion of the native *tatC* gene.

[7] To obtain JOE2475, pJC368 was mated into NA1000 for integration at the *CC2002* locus, and subsequently pJC366 was integrated at the *xylX* locus, also introduced by mating. The resulting strain was counter-selected with sucrose in the presence of xylose, and sucrose-insensitive isolates were screened for the deletion of the native *tatB* gene.

[8] To obtain PBP2 depletion strains JOE7298 and JOE7302, pJC774 was first electroporated into NA1000 for integration at the *xylX* locus to generate strains JOE7294 and JOE7295, and subsequently pTC358 was electroporated for integration at the *pbp2* locus. The resulting strains (including JOE7296) were counter-selected with sucrose in the presence of xylose, and sucrose-insensitive isolates were screened for the loss of pTC358 and deletion within the native *pbp2* gene. JOE7298 – JOE7302 were isolates with the deletion, while JOE7303 was an isolate that retained the native *pbp2* gene.

**TABLE S2.** Plasmids used for this study.

| Plasmids | Relevant genetic markers, features, and/or description [1] | Source / Notes |
|---|---|---|
| pCM62 | RK2-derived broad-host-range vector with *E. coli lac* promoter preceding polylinker; $Tc^R$ | (Marx & Lidstrom, 2001) |
| pEJ216 | pJS14-$_{ss}$*torA-dimer2*; $Cm^R$ | (Judd *et al.*, 2005) [2] |
| pJS14 | pBBR1-based broad-host-range vector; $Cm^R$ | (Jenal & Fuchs, 1998) |
| pMT69 | for integration into *C. crescentus xylX* locus; $Sp^R$ | (Thanbichler and Shapiro, 2006) |
| pNPTS138 | for allelic replacement during two-step selection; *sacB*, $Km^R$ | M.R.K. Alley |
| pPR9TT | RK2-derived vector for translational fusions to *lacZ*; $Ap^R$, $Cm^R$ | (Santos *et al.*, 2001) |
| pVCHYN-5 | for generating N-terminal protein fusion to mCherry, encoded at the *C. crescentus vanA* locus; $Tc^R$ | (Thanbichler *et al.*, 2007) |
| pXGFP4-C1 | for generating N- terminal protein fusion to GFP, encoded at the *C. crescentus xylX* locus; $Km^R$ | M.R.K. Alley |
| pX-mCh-*mreB* | plasmid derived from pXGFP4-C1, for integrating into *C. crescentus xylX* locus and placing *mCherry-mreB* under $P_{xyl}$ control | (Dye *et al.*, 2005) |
| pX-*mreC-mCherry* | plasmid derived from pXGFP4-C1, for integrating into *C. crescentus xylX* locus and placing *mreC-mCherry* under $P_{xyl}$ control | (Dye *et al.*, 2005) |
| pXGFP4C1-*pbp2* | plasmid derived from pXGFP4-C1, for integrating into *C. crescentus xylX* locus and placing *gfp-pbp2* under $P_{xyl}$ control | (Dye *et al.*, 2005) |
| pJC305 | pNPTS138-Δ*CC2001*; for in-frame deletion of *CC2001* | This study [3] |
| pJC324 | pCM62-*rrnB* T1T2 (transcriptional terminator); $Tc^R$ | (Chen *et al.*, 2006) |
| pJC326 | pJC324-*lacZ*; $Tc^R$ | (Chen *et al.*, 2006) |
| pJC349 | pNPTS138-Δ*CC2001*::Ω*aacC4*; for allelic replacement of *CC2001* | This study [4] |
| pJC351 | pMT69-*CC2001* ($P_{xyl}$-*CC2001* integration); $Sp^R$ | This study [5] |
| pJC363 | pPR9TT-$P_{xyl}$-*CC2001*; $Cm^R$ | This study [6] |
| pJC366 | pMT69-*CC2002* ($P_{xyl}$-*CC2002* integration); $Sp^R$ | This study [7] |
| pJC368 | pNPTS138-Δ*CC2002*; for in-frame deletion of *CC2002* | This study [8] |
| pJC372 | pNPTS138-Δ*CC2003*::*aacC4*; for allelic replacement of *CC2003* | This study [9] |
| pJC378 | pCM62-*CC2003-CC2001-lacZ* (derived from pJC326); $Tc^R$ | This study [10] |
| pJC386 | pPR9TT with additional restriction sites; $Cm^R$ | This study [11] |
| pJC392 | pCM62-*CC2003-CC2002-lacZ* (derived from pJC326); $Tc^R$ | This study [12] |
| pJC396 | pPR9TT-$P_{xyl}$-*CC2002* (derived from pJC386); $Cm^R$ | This study [13] |
| pJC397 | pPR9TT-$P_{xyl}$-*CC2003* (derived from pJC386); $Cm^R$ | This study [14] |
| pJC399 | pMT69-*CC2003* ($P_{xyl}$-*CC2003* integration); $Sp^R$ | This study [15] |
| pJC402 | pNPTS138-Δ*CC2003-CC2001*::*aacC4*; for allelic replacement of *tatABC* | This study [16] |
| pJC424 | pVan-*mCherry-mreB*; $Tc^R$ | This study [17] |
| pJC427 | pVan-*mCherry-pbp2*; $Tc^R$ | This study [18] |
| pJC774 | pXyl-*CC1546* ($P_{xyl}$-*pbp2* integration); $Gm^R$ | This study [19] |
| pTC343 | pVan-*mCherry-pbp1a*; $Tc^R$ | This study [20] |
| pTC358 | pNPTS138-Δ*pbp2*; for in-frame deletion of PBP2's enzymatic region | This study [21] |

[1] Ap, Cm, Km, Sp, Tc: abbreviations for ampicillin, chloramphenicol, kanamycin, spectinomycin, (oxy)tetracycline.

[2] Plasmid pEJ216 was originally published as expressing ssTorA-tdimer2 from a xylose-regulated promoter ($P_{xyl}$) on pJS14. However, more recent whole-plasmid sequencing indicated that pEJ216 actually expresses ssTorA-dimer2 from the constitutive *E. coli lac* promoter ($P_{lac}$). ssTorA represents the first 40 amino acids of *E. coli* trimethylamine *N*-oxide reductase, including its canonical Tat signal sequence (Cristóbal *et al.*, 1999). The fluorescent protein dimer2 is a derivative of DsRed, while tdimer2 contains two linked, tandem copies of dimer2 (Campbell *et al.*, 2002).

[3] For construction of pJC305, the region upstream of *CC2001* was amplified with primers CC2001 -585F and CC2001 20R and digested with EcoRI and BamHI, and the region downstream of *CC2001* was amplified with primers CC2001 885F and CC2001 +575R and digested with BamHI and HindIII; the two resulting fragments were inserted into pNTPS138 digested with EcoRI and HindIII, fusing the first seven codons and last six codons (including the stop codon) of *CC2001* in-frame.

[4] For construction of pJC349, pHP45Ωaac4 (Blondelet-Rouault *et al.*, 1997) was digested with BamHI to release the apramycin resistance cassette, which was subsequently inserted into the BamHI site of pJC305; the *aacC4* gene is in the reverse orientation relative to the replaced *CC2001* gene.

[5] For construction of pJC351, *CC2001* was amplified with primers CC2001 F and CC2001 916R, digested with EcoRI and HindIII, and inserted into pJC300 (Chen *et al.*, 2005) digested with the same enzymes to yield pJC342. A fragment containing *CC2001* was then obtained from pJC342 by digesting with XhoI and AflII and inserted into pMT69 digested with the same enzymes to yield pJC351.

[6] For construction of pJC363, $P_{xyl}$-*CC2001* was amplified from pJC351 with primers Pxyl -506F and CC2001 +14R, digested with BglII and HindIII, and inserted into pPR9TT digested with the same enzymes.

[7] For construction of pJC366, *CC2002* was amplified with primers CC2002 F and CC2002 +11R, digested with EcoRI and HindIII, and inserted into pJC351 digested with the same enzymes to replace *CC2001*.

[8] For construction of pJC368, the region upstream of *CC2002* was amplified with primers CC2002 -553F and CC2002 23R and digested with HindIII and BamHI, and the region downstream of *CC2002* was amplified with primers CC2002 588F and CC2002 +538R and digested with BamHI and EcoRI; the two resulting fragments were inserted into pNTPS138 digested with EcoRI and HindIII, fusing the first eight and last five codons of *CC2002* in-frame.

[9] For construction of pJC372, the region upstream of *CC2003* was amplified with primers CC2003 -517F and CC2003 29R and digested with HindIII and BamHI, and the region downstream of *CC2003* was amplified with primers CC2003 204F and CC2002 +565R and digested with BamHI and EcoRI; the two resulting fragments were inserted into pNTPS138 digested with EcoRI and HindIII, fusing the first ten and last seven codons of *CC2003* to yield pJC369. The *aacC4* gene was amplified from pHP45Ωaac4 with primers aacC4 F BamHI and aacC4 R BamHI, digested with BamHI, and inserted into the BamHI site of pJC369 for in-frame fusion to the start and stop codons of *CC2003*, yielding pJC372.

[10] For construction of pJC378, *tatABC* was amplified with primers CC2004 626F and CC2001 916R, digested with BglII and HindIII, and inserted into pJC326 digested with the same enzymes.

[11] For construction of pJC386, a linker containing AflII, NheI, and SphI sites was inserted into pPR9TT between the XmaI and BamHI sites.

[12] For construction of pJC392, *tatAB* was amplified with primers CC2004 626F and CC2002 +11R, digested with BglII and HindIII, and inserted into pJC378 digested with the same enzymes to replace *tatABC*.

[13] For construction of pJC396, a PstI-SphI fragment from pJC366 containing *CC2002* was inserted into the same sites of pJC386.

[14] For construction of pJC397, *CC2003* was amplified with primers CC2004 626F and CC2002 2R, digested with BglII and HindIII, and inserted into pJC378 digested with the same enzymes, replacing *tatABC* and yielding pJC393. A BglII-SphI fragment from pJC393 was then inserted into the same sites of pJC386 to yield pJC397.

[15] For construction of pJC399, *CC2003* was amplified with primers CC2003 -28F and CC2003 +1R, digested with EcoRI and HindIII, and inserted into pJC366 digested with the same enzymes to replace *CC2002*.

[16] For construction of pJC402, a SpeI-BamHI fragment from pJC369 (precursor of pJC372) containing the region upstream of *CC2003* was inserted into pJC305 cut with NheI and BamHI, thus replacing the region upstream of *CC2001*, fusing the first ten codons of *CC2003* in-frame to the BamHI site and the last five codons of *CC2001*, and yielding pJC400. A BamHI fragment containing the *aacC4* gene from pJC372 was then inserted into the BamHI site of pJC400, between the region upstream of *CC2003* and the region downstream of *CC2001*, to generate pJC402.

[17] For construction of pJC424, an NdeI-XbaI fragment containing *mCherry-mreB* from pX-mCh-*mreB* was inserted into pVCHYN-5 digested with NdeI and NheI.

[18] For construction of pJC427, a KpnI-XbaI fragment containing *pbp2* from pXGFP4C1-*pbp2* was inserted into pVCHYN-5 digested with KpnI and NheI.

[19] For construction of pJC774, *CC1546* was amplified with primers MP004F and MP004R and inserted into pXYFPC-4 (Thanbichler *et al.*, 2007) digested with NdeI and NheI, via Gibson assembly.

[20] For construction of pTC343, *CC1516* was amplified with primers TC695F and TC696R and inserted into pVCHYN-5 digested with KpnI and NheI, via Gibson assembly.

[21] For construction of pTC358, a region upstream and overlapping the 5' portion of *CC1546* was amplified with primers TC739F and TC740R, and a region downstream and overlapping the 3' portion of *CC1546* was amplified with primers TC741F and TC742R; the two resulting fragments were inserted into pNTPS138, digested with SpeI and EcoRI, via Gibson assembly.

**TABLE S3.** Primers used for this study.

| Primer | Sequence (5' to 3') |
|---|---|
| CC2001 -585F | tgcgaattccggcggcacagaacttctca |
| CC2001 20R | gtaggatccgtgtccgatggctttactca |
| CC2001 885F | gttggatccgacctggtcgtttcgtaag |
| CC2001 +575R | cctaagcttcctcggtcgagatccagaac |
| CC2001 F | caggaattcgtgagtaaagccatcggacac |
| CC2001 916R | ctcaagcttagaccgcccttgccttacga |
| Pxyl -506F | ccgagatctaggtcttccaccagccacag |
| CC2001 +14R | gccaagcttagaccgcccatgccttacga |
| CC2002 F | cgcgaattcatgcttcctgatatcggcggcaca |
| CC2002 +11R | gcgaagcttgatggctttactcacgagac |
| CC2002 -553F | ttcaagcttccggacccgttgggtctgag |
| CC2002 23R | gatggatcctgtgccgccgatatcaggaa |
| CC2002 588F | cgaggatccgacatcgtctcgtgagtaaa |
| CC2002 +538R | gtcgaattcctcagcgagaaccacagaac |
| CC2003 -517F | gagaagcttgccatcgtcgcctatcacca |
| CC2003 29R | gaaggatcccacccagtgaatccaactca |
| CC2003 204F | gcaggatccgaagagcttcgcaagtcgtaa |
| CC2003 +565R | tgcgaattccttgaccggctcgaccaaga |
| aacC4 F BamHI | cgtggatcccaatacgaatggcgaaaag |
| aacC4 R BamHI | ctcggatcctgagctcagccaatcgactg |
| CC2004 626F | cacagatctattcgcgcaggatttcctgg |
| CC2002 2R | atcaagcttcatggacgcgccttagagc |
| CC2003 -28F | ggtgaattccgccgctgtcgcaaggagca |
| CC2003 +1R | ctgaagcttacgacttgcgaagctct |
| MP004F | tcggcgcttcagacgctcgagttttggggagacgaccatatgagcgaaccgtccatcttc |
| MP004R | gaatggccgctctagaactagtggatcccccgggctgcagctagctcatgtctggcctcc |
| TC695F | gcgccttaattaatatgcatggtaccatgtctgatcctaccgaccct |
| TC696R | aactagtggatcccccgggctgcagctagctcaggggcgcggc |
| TC739F | gtgcaattgaagccggctggcgccaagcttatgagcgaaccgtccatcttctttttcg |
| TC740R | ccggatccttcaggcggttctggatatcggcg |
| TC741F | tatccagaaccgcctgaaggatccggagatccgcg |
| TC742R | atccggagacgcgtcacggccgaagctagcgtcgtgcagaacgaacatcacaaag |

### Supplemental Figure Legends

**Figure S1.** Colonies and cell morphology of *tat* and *pbp2* strains. (**A**) Plasmid loss and colony-sectoring assay. Strains with deletions of *tat* genes are unable to lose their complementing plasmids. NA1000- or CB15-derived strains were constructed with wild-type alleles or deletions of *tat* genes (Δ) on the chromosome while carrying the corresponding *tat* genes and the *E. coli lacZ* gene on complementing plasmids. They were streaked onto PYE plates containing X-Gal and incubated at 30°C. Resultant colonies were photographed, sometimes after plates were refrigerated for several days to enhance the blue color. Images shown here demonstrate the variations in colony and background colors due to factors such as incubation period and camera lighting, but all indicate that deletion strains do not produce white colonies because they are unable to lose the *lacZ* gene, linked to the complementing *tat* allele. Strains used were constructed from NA1000 [JOE2670 and JOE2692 (*tatA*), JOE2713 and JOE2712 (*tatB*), JOE2402 and JOE2406 (*tatC*), JOE2856 and JOE2877 (*tatABC*)] or CB15 [JOE2674 and JOE2696 (*tatA*), JOE2749 and JOE2748 (*tatB*), JOE2403 and JOE2408 (*tatC*), JOE2858 and JOE2890 (*tatABC*)]. Images are representative of two biological replicates. (**B**) Cell morphology of wild-type strains expressing *tat* or *pbp2* from the *xylX* promoter ($P_{xyl}$). Phase contrast images were acquired after expression strains were grown under inducing (xylose) or non-inducing (glucose) conditions for 22-24 hours (n = 3 biological replicates). Strains used were JOE2719, JOE2758, JOE2332, and JOE7294. Scale bar, 5 μm.

**Figure S2.** PBP2 is essential for colony formation in *Caulobacter*. (**A**) PBP2 is required for robust colony formation. Strains were streaked onto PYE plates containing 0.1% xylose, 0.1% glucose, or 5 μg/mL gentamicin (Gm) + 0.2% glucose. Relevant genotypes of strains in each sector are shown in the schematic on the left. Strains used were JOE2357, JOE7295, JOE7296, JOE7301, JOE7302, and JOE7303. JOE7296 (*pbp2*::* $P_{xyl}$-*pbp2*) is derived from JOE7295 and carries the allelic replacement plasmid pTC358 integrated into the *pbp2* locus. Strains with deletion (JOE7301 and JOE7302) or wild-type (JOE7303) allele of *pbp2* were derived from JOE7296 following sucrose counter-selection. Images represent results from three or more independent replicates, each done on a separate day. (**B**) *mCherry-pbp2* can complement null mutation in *pbp2*. Strains with or without the *pbp2*$\Delta_{TP}$ mutation and expressing mCherry or mCherry-PBP2 from the vanillate-inducible promoter ($P_{van}$) were streaked onto PYE plates containing 0.1% xylose + 1 μg/mL oxytetracycline (Tc) or 0.1% glucose to induce or repress, respectively, $P_{xyl}$-*pbp2*. Basal expression of mCherry-PBP2 from the vanillate promoter is sufficient for complementation. Schematic on the left indicate relevant genotypes of strains in each quadrant. Strains used were JOE7324, JOE7326, JOE7328, and JOE7329. Image shown represents results from three or more independent trials.

**Figure S3.** Midcell localization of PBP2 and PBP1a in Tat depletion strains upon osmotic upshift. Cultures were induced for four hours with vanillate to express (**A**, **B**, **C**, **D**) mCherry-PBP2 or (**E**, **F**) mCherry-PBP1a. (**A**, **B**) TatB depletion strain (Δ*tatB* $P_{xyl}$-*tatB*) was grown in the presence of xylose or glucose for four hours to express or repress *tatB*, respectively, while (**C**, **D**) TatC depletion strain (Δ*tatC* $P_{xyl}$-*tatC*) was first grown for 14 – 16 hours with glucose and then resuspended in PYE medium containing xylose or glucose for four additional hours of growth. (**E**, **F**) WT and TatA depletion strains expressing mCherry-PBP1a were grown similarly as those in Figure 2, in the absence or presence of xylose or glucose for four hours. Cells were subjected to osmotic upshift from being transferred from PYE medium to an M2 agarose pad for microscopy. (**A**, **C**, **E**) Representative phase contrast (top) and fluorescence (middle) images are shown with corresponding population-level demographs (bottom). Demographs depict localization of normalized fluorescence along the medial axis (the cell length), with cells arranged by length and lighter colors indicating brighter fluorescence. (**B**, **D**, **F**) Medial profiles (left panels) represent normalized fluorescence intensities along normalized cell length, as illustrated by the schematic of a model cell to the left of part (B). Colored lines indicate averages, while shaded areas indicate 95% confidence intervals. Area under curve was calculated for each medial profile and shown at the population level as scatter plots (right panels), with horizontal bars indicating means and standard deviations. (**A**, **B**) Midcell localization of mCherry-PBP2 is ineffective in the TatB depletion strain, regardless of TatB expression; n = 1146 (WT), 1853 (Δ*tatB* + Xyl), 1611 (Δ*tatB* + Glu). (**C**, **D**) Midcell localization of mCherry-PBP2 depends on TatC; n = 1568 (WT), 1683 (Δ*tatC* + Xyl), 2126 (Δ*tatC* + Glu). (**E**, **F**) Midcell localization of mCherry-PBP1a is not affected by TatA depletion; n = 1415 (WT), 877 (Δ*tatA* + Xyl), 991 (Δ*tatA* + Glu). Scale bar, 5 µm. **, $p < 0.01$; ****, $p < 0.0001$; based on two-tailed *t*-test.

**Figure S4.** Localization of MreB and PBP1a without osmotic upshift. WT and TatA depletion strains were induced with vanillate to express (**A**, **B**) mCherry-MreB or (**C**, **D**) mCherry-PBP1a and grown in the absence or presence of xylose or glucose, as described in Figure 2, and transferred from PYE medium to PYE agarose pads for microscopy, without being subject to osmotic shock. (**A**, **C**) Representative phase contrast (top) and fluorescence (bottom) images are shown. Scale bar, 5 µm. (**B**, **D**) Transverse profiles (left panels) represent normalized fluorescence intensities along the cell width (minor axis), as illustrated by the schematic of a model cell on the left. Colored lines indicate averages, while shaded areas indicate 95% confidence intervals. Normalized fluorescence at the midpoint of each transverse profile is shown at the population level as scatter plots (right panels), with horizontal bars indicating means and standard deviations. 400 cells were measured for each mCherry-MreB population, while 100 cells were measured for each mCherry-PBP1a population. No significant differences were observed among populations with the same fluorescence fusion, based on two-tailed *t*-tests.
